# Molecular de-extinction of ancient antimicrobial peptides enabled by machine learning

**DOI:** 10.1101/2022.11.15.516443

**Authors:** Jacqueline R. M. A. Maasch, Marcelo D. T. Torres, Marcelo C. R. Melo, Cesar de la Fuente-Nunez

## Abstract

Molecular de-extinction could offer new avenues for drug discovery by reintroducing bioactive molecules that are no longer encoded by extant organisms. To prospect for antimicrobial peptides encrypted as subsequences of extinct and extant human proteins, we introduce the panCleave random forest model for proteome-wide cleavage site prediction. Our model outperformed multiple protease-specific cleavage site classifiers for three modern human caspases, despite its pan-protease design. Antimicrobial activity was observed *in vitro* for modern and archaic protein fragments identified with panCleave. Lead peptides were tested for mechanism of action, resistance to proteolysis, and anti-infective efficacy in two pre-clinical mouse models. These results suggest that machine learning-based encrypted peptide prospection can identify stable, nontoxic antimicrobial peptides. Moreover, we establish molecular de-extinction through paleoproteome mining as a framework for antibacterial drug discovery.

**Highlights:**

1. Machine learning guides bioinspired prospection for encrypted antimicrobial peptides.
2. Modern and extinct human proteins harbor antimicrobial subsequences.
3. Ancient encrypted peptides display *in vitro* and *in vivo* activity with low host toxicity.
4. Paleoproteome mining offers a new framework for antibiotic discovery.

## Introduction

The idea of reintroducing extinct organisms into extant environments has captured the public and scientific imagination, raising profound ethical and ecological questions (*1*). Here, we introduce molecular de-extinction as an antibiotic discovery framework. Molecular de-extinction is the resurrection of extinct molecules of life: nucleic acids, proteins, and other compounds no longer encoded by living organisms. While the societal benefit of organismal de-extinction is still unknown and contentious, technical challenges like incomplete genomic coverage remain significant (*1, 2*). By synthesizing only isolated compounds, molecular de-extinction circumvents many of the ethical and technical problems posed by whole-organism de-extinction. Molecular de-extinction is motivated by the hypothesis that molecules that conferred benefits to extinct organisms could be beneficial in the current global environment. Such molecules could be of biomedical or economic utility by bolstering defenses against future challenges that resemble stressors from environments past, including climate change or infectious disease outbreaks. The present work proposes molecular de-extinction as a drug discovery framework for expanding the therapeutic search space through paleoproteome mining.

The global antibiotic resistance crisis, the threat of emerging pathogens, and the overuse of traditional antibiotic scaffolds necessitate new, computer-aided drug development paradigms (*3*). Protein informatics is fertile ground for antibiotic discovery, as many peptides are known to modulate the host immune system, disrupt bacterial cell membranes, suppress biofilms, and promote wound healing (*4*). Furthermore, 20^*n*^ variants exist per *n*-length canonical amino acid sequence, presenting an enormous combinatorial space from which to select peptides with targeted activity. Antimicrobial peptides (AMPs) are an ancient class of host defense molecule found across the domains of life, representing an essential facet of innate immunity throughout evolution. Some AMPs have demonstrated collateral sensitivity in antibiotic-resistant bacteria and a low propensity to induce resistance (*4, 5*). The human cryptome is a subset of the proteome known to harbor AMPs that are released from precursor proteins by both host and pathogen proteases (*6, 7*). These bioactive encrypted peptides can serve as natural templates for new antibiotics and for bioinspired engineered therapies (*8*).

To mine extinct and extant human proteomes for potential encrypted peptides, we present the panCleave Python pipeline (https://gitlab.com/machine-biology-group-public/pancleave). This open-source machine learning (ML) tool leverages a pan-protease cleavage site classifier to perform computational proteolysis: the *in silico* digestion of human proteins into peptide fragments (Figs. 1, S1). We experimentally validate panCleave for the prospection of encrypted AMPs in modern human secreted proteins and in the archaic proteomes of our closest extinct relatives, Neanderthals and Denisovans (Fig. 1). Using panCleave, we discovered new peptides encrypted within known precursor protein groups and rediscovered a known encrypted AMP. By discovering novel AMPs through computational paleoproteome mining, this work offers a proof-of-concept for molecular de-extinction as an antibiotic discovery framework. Furthermore, this study introduces the first known antimicrobial subsequences encrypted in archaic human proteins.

**Fig. 1.**
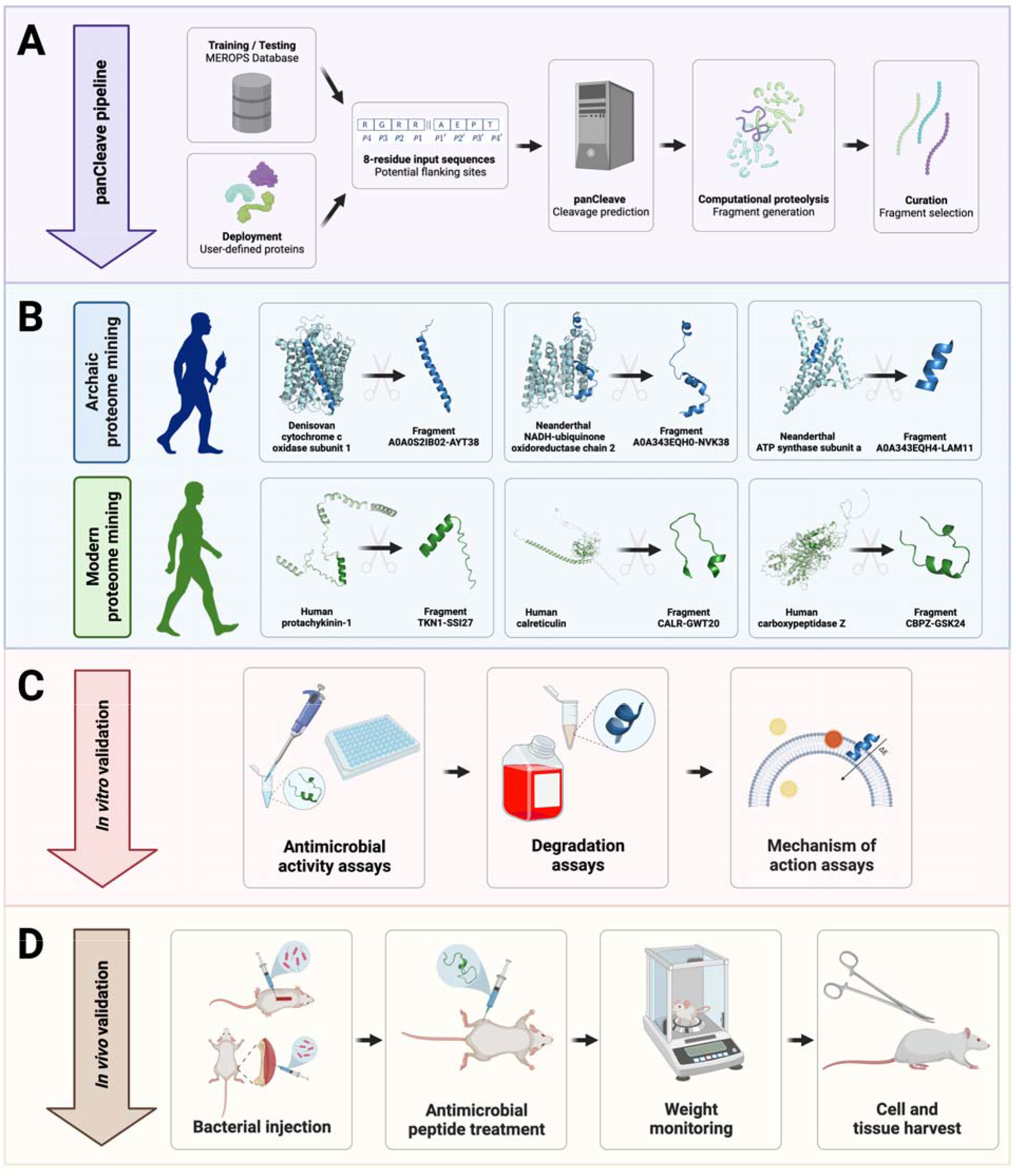
Computational-experimental framework for molecular de-extinction of antimicrobial peptides. Panel (**A**) demonstrates the computational proteolysis pipeline, where user-defined proteins are processed into 8-residue subsequences that are classified as cleavage and non-cleavage sites. Input proteins are then tokenized at predicted cleavage sites, and the resulting fragments can be filtered by user-defined curation methods. Curation methods can include machine learning-based activity prediction, human expert curation, or other methods. Successes in archaic and modern proteome mining are visualized in panel (**B**), where precursors were computationally digested to reveal encrypted antimicrobial subsequences. The pipeline concludes with *in vitro* (**C**) and *in vivo* (**D**) experimental validation of fragment bioactivity, including proteolytic degradation assays, MoA assays, and mouse weight monitoring as a proxy for host toxicity. Figure created with BioRender.com and the PyMOL Molecular Graphics System, Version 2.1 Schrödinger, LLC.

## Results

### Computational proteolysis pipeline

The panCleave Python pipeline (Figs. 1, S1) is a protein informatics tool that uses ML for computational proteolysis: the *in silico* fragmentation of human protein sequences into peptides. The development of this predictive tool was motivated by the hypothesis that protease-agnostic cleavage site prediction could facilitate biologically inspired prospection for encrypted host defense peptides. Prior cleavage site classifiers are specialized models that predict cleavage activity for only a subset of human proteases (*9–20*). A pan-protease design facilitates proteome-wide searches, circumventing the need to hypothesize protease-substrate relationships. To our knowledge, the panCleave random forest is the first cleavage site classifier trained on all human protease substrates in the MEROPS Peptidase Database (*21*). Substrate amino acid frequencies, length distributions, protease representation, and precursor protein functions for all training and testing data are characterized in Figs. S2–S6. Source code, training data, and testing data are available on GitLab (https://gitlab.com/machine-biology-group-public/pancleave).

The performance of the panCleave random forest can be quantified on an aggregated, protease-agnostic level and a disaggregated, protease-specific level. On the complete independent test set comprising substrates from 182 proteases (*n* = 9,927), panCleave achieved an overall accuracy of 73.3%. Thresholding by estimated probability of binary class membership (*i.e*., probability that a subsequence is a cleavage site or non-cleavage site) indicates increasing accuracy with increasing estimated probability: panCleave achieved 81.9% accuracy for predictions of 60% estimated probability or greater (62.8% of test set predictions) and a maximum accuracy of 96.6% for predictions of 90% estimated probability or greater (2.1% of predictions) (Fig. 2c). The random forest probability estimate is useful for providing a degree of confidence in a predicted class membership (*22*). The area under the receiver operating characteristic curve was 80.8% and the average precision was 80.3% (Fig. 2a,b). Negative predictive value, positive predictive value, sensitivity, and specificity were 73.2%, 73.5%, 73.0%, and 73.6%, respectively.

**Fig. 2.**
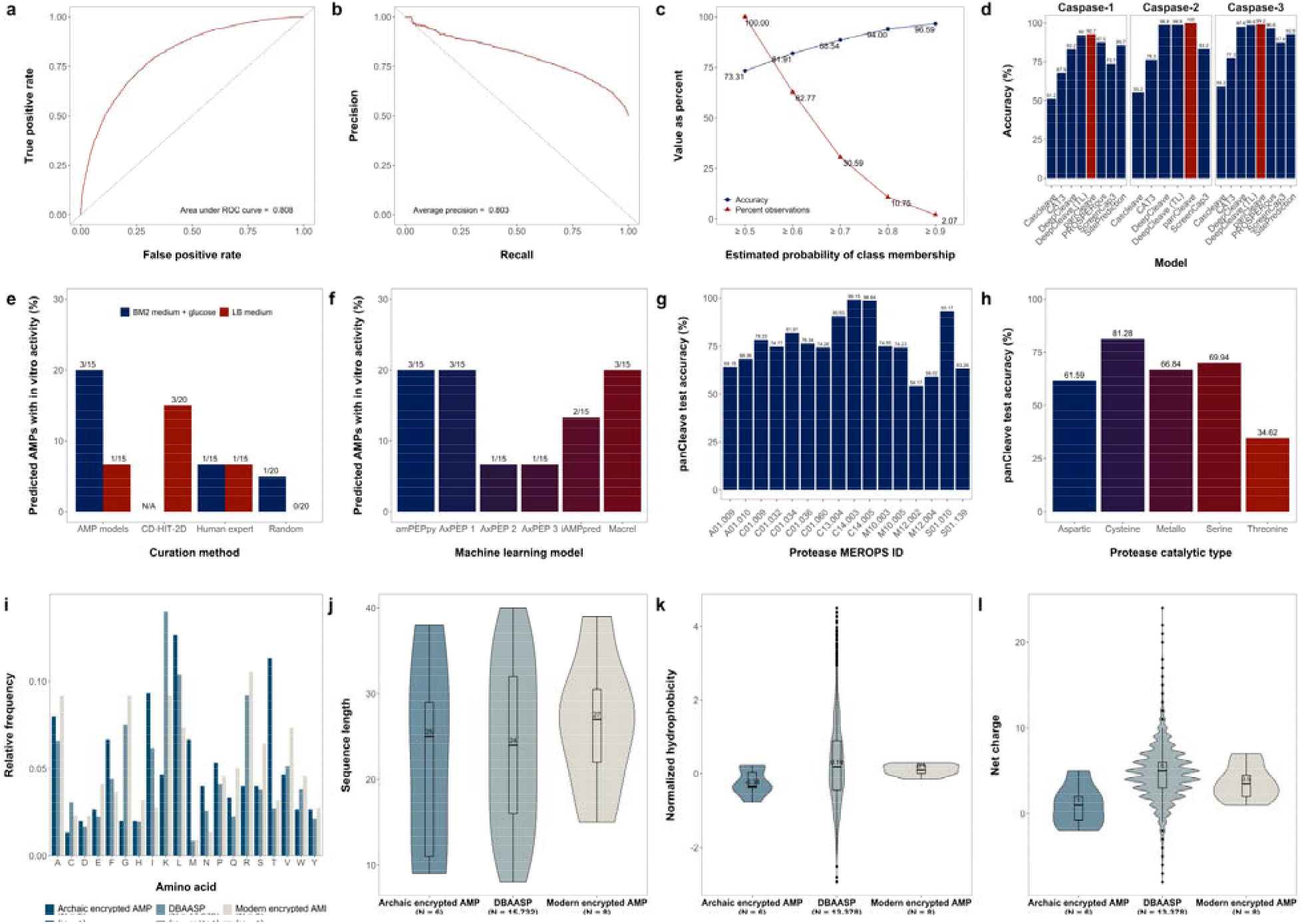
Model performance and antimicrobial peptide data distributions. Panels describe panCleave random forest performance evaluation (**a-h**) and physicochemical distributions for positive hits (**i–l**). Optimized panCleave random forest performance is reported for independent test data (*n* = 9,927): (**a**) the receiver operating characteristic curve; (**b**) precision-recall curve; (**c**) accuracy-probability threshold tradeoff curves; (**d**) accuracy of panCleave relative to preexisting models for three caspases; (**e**) positive hit rate by fragment curation method; (**f**) positive hit rate by antimicrobial activity classifier; (**g**) panCleave test accuracy for proteases with at least 100 test observations; and (**h**) panCleave test accuracy by protease catalytic type. Panels **i–l** compare amino acid frequency (**i**), fragment length (**j**), normalized hydrophobicity (**k**), and net charge distributions (**l**) for modern encrypted AMPs, archaic encrypted AMPs, and AMPs reported in DBAASP (*23*). Hydrophobicity scores employ the Eisenberg and Weiss scale (*25*). Note that DBAASP data were restricted to fragments of length 8–40 residues for length, hydrophobicity, and charge distributions, with null values excluded. DBAASP amino acid frequencies were computed by excluding noncanonical residues.

When disaggregating model accuracy by protease, panCleave performance ranged widely (Fig. 2g,h; Tables S1–S4). Among proteases with at least 100 test set observations, panCleave achieved greater than 80% accuracy on caspase-3 (C14.003; 99.2%), caspase-6 (C14.005; 98.6%), granzyme B (S01.010; 93.2%), legumain (C13.004; 90.6%), and cathepsin S (C01.034; 81.9%) (Fig. 2g; Table S1). Among protease clans, panCleave achieved greater than 70% accuracy on endopeptidase clan CD (type protease caspase-1 [C14.001]; 93.9%), endopeptidase/exopeptidase clan SB (type protease subtilisin Carlsberg [S08.001]; 88.6%), cysteine protease clan CA (type protease papain [C01.001]; 74.1%), and endopeptidase clan PA (type protease chymotrypsin A [S01.001]; 70.6%) (Table S3). The average accuracy was greatest for cysteine catalytic types (81.3%; 1858/2286 observations predicted correctly) and lowest for threonine catalytic types (34.6%; 18/52) (Fig. 2h).

When compared to pre-existing protease-specific models, panCleave outperformed for caspase-2 (C14.006; 100.0%), caspase-3 (C14.003; 99.15%), and caspase-1 (C14.001; 92.68%) (Figure 2d; Table S4). However, pre-existing models outperformed for multiple matrix metallopeptidases (Table S4). While the pan-protease design of panCleave does not preclude the possibility of high or state-of-the-art accuracy for specific proteases, the use of panCleave for protease-specific applications should be guided by the reported disaggregated accuracies (Fig. 2d,g,h; Tables S1– S4).

### Modern encrypted peptides display antimicrobial activity in vitro

Eight of 80 (10.0%) modern secreted protein fragments were active against one or more pathogens in at least one of the conditions tested (Fig. 3; Tables S5, S6). Importantly, none of the tested sequences have yet been reported as AMPs or as AMP subsequences in the Database of Antimicrobial Activity and Structure of Peptides (DBAASP) (*23*).

**Fig. 3.**
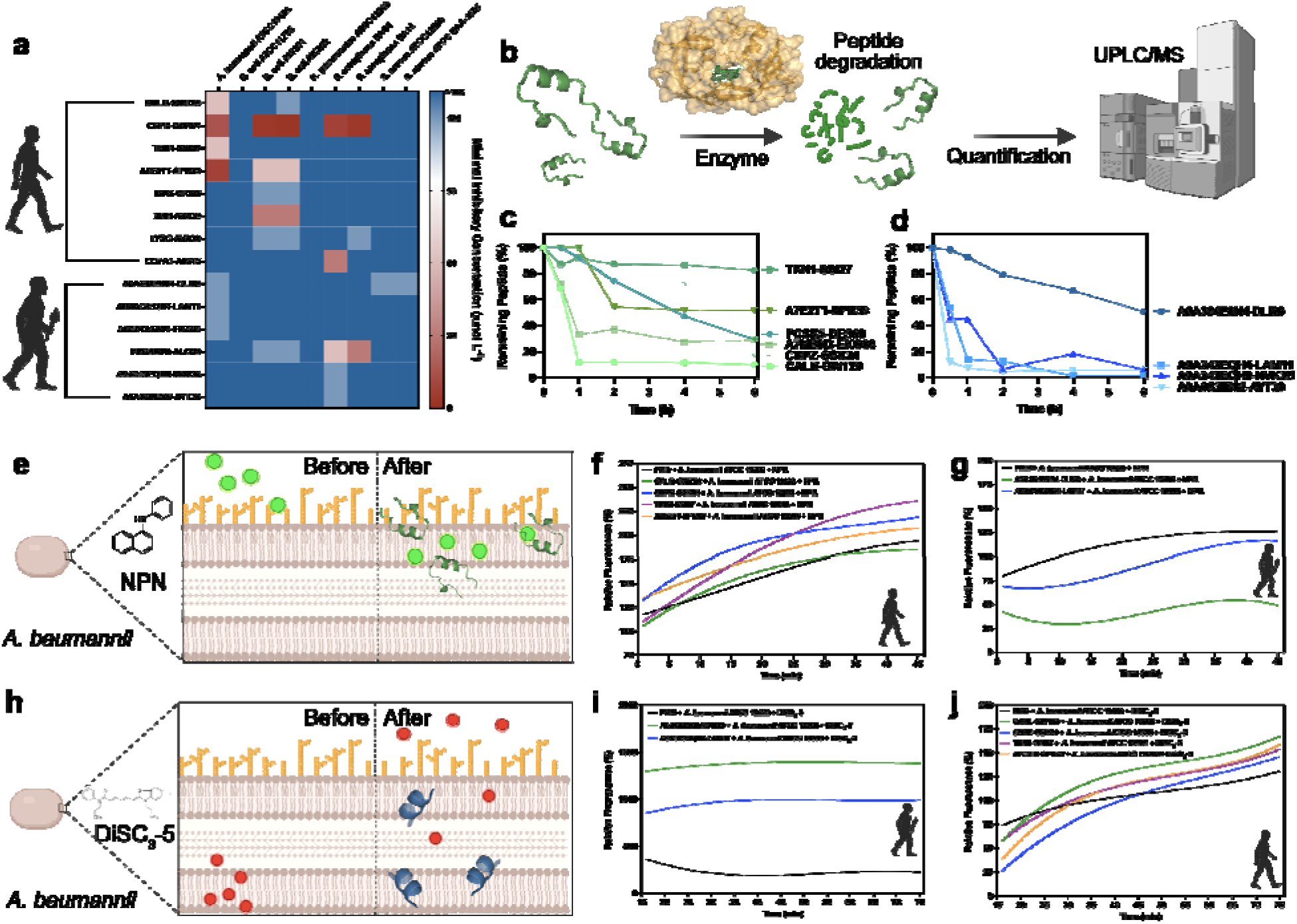
Antimicrobial activity, resistance to enzymatic degradation, and mechanism of action of modern and archaic encrypted peptides. **(a)** Antimicrobial activity of the encrypted peptides. Briefly, a fix number of 10^6^ bacterial cells per mL^−1^ was used in all the experiments. The modern and archaic encrypted peptides were two-fold serially diluted ranging from 128 to 2 μmol L^−1^ in a 96-wells plate and incubated at 37□°C for one day. After the exposure period, the absorbance of each well was measured at 600 nm. Untreated solutions were used as controls and minimal concentration values for complete inhibition were presented as a heat map of antimicrobial activities (μmol L^−1^) against nine pathogenic bacterial strains. All the assays were performed in three independent replicates and the heat map shows the mode obtained within the two-fold dilutions concentration range studied. **(b)** Schematic of the resistance to enzymatic degradation experiment, where peptides were exposed for a total period of six hours to fetal bovine serum that contains several active proteases. Aliquots of the resulting solution were analyzed by ultra-performance liquid chromatography coupled to mass spectrometry (UPLC/MS). **(c)** Modern and **(d)** archaic peptides had different degradation behaviors. In summary, archaic peptides are more resistant to enzymatic degradation than modern peptides. Experiments were performed in two independent replicates. **(e)** Schematic showing the behavior of 1-(N-phenylamino)naphthalene (NPN) the fluorescent probe used to indicate membrane permeabilization caused by the encrypted peptides. **(f)** Modern and **(g)** archaic encrypted peptides fluorescence values relative to the untreated control showing that modern peptides are more efficient to permeabilize the outer membrane of *A. baumannii* cells than polymyxin **B** (PMB) and archaic encrypted peptides. **(h)** Schematic of how 3,3,-dipropylthiadicarbocyanine iodide [DiSC_3_-(5)], a hydrophobic fluorescent probe, was used to indicate membrane depolarization caused by the encrypted peptides. **(i)** Modern and **(j)** archaic encrypted peptides fluorescence values relative to the untreated control showing that archaic peptides are much stronger depolarizers of the cytoplasmic membrane of *A. baumannii* cells than polymyxin B (PMB) and modern encrypted peptides. Experiments were performed in three independent replicates. Figure created with BioRender.com and the PyMOL Molecular Graphics System, Version 2.1 Schrödinger, LLC.

The encrypted peptide CBPZ-GSK24 from carboxypeptidase Z (UniProt ID: CBPZ_HUMAN) demonstrated the strongest and most broad-spectrum antimicrobial activity *in vitro*, inhibiting *Pseudomonas aeruginosa* PA01 (8 μmol L^−1^), *Pseudomonas aeruginosa* PA14 (4 μmol L^−1^), *Escherichia coli* AIC221 (4 μmol L^−1^), *Escherichia coli* AIC222 (2 μmol L^−1^), and *Acinetobacter baumannii* ATCC19606 (16 μmol L^−1^). Fragment A7E2T1-SPR29 of uncharacterized protein A7E2T1_HUMAN also displayed broad-spectrum activity against *E. coli* AIC221 (64 μmol L^−1^), *E. coli* AIC222 (64 μmol L^−1^), and *A. baumannii* ATCC19606 (8 μmol L^−1^). CALR-GWT20, encrypted in calreticulin (UniProt ID: CALR_HUMAN), displayed antimicrobial activity against colistin-resistant *E. coli* AIC222 at 128 μmol L^−1^ and *A. baumannii* ATCC19606 at 64 μmol L^−1^. Fragment XDH-AVA32, a subsequence of xanthine dehydrogenase/oxidase (UniProt ID: XDH_HUMAN), was active at 32 μmol L^−1^ against both *E. coli* AIC221 and AIC222 strains. ISK5-GKI32, part of the serine protease inhibitor kazal-type 5 (UniProt ID: ISK5_HUMAN), was also active at 128 μmol L^−1^ against both *E. coli* strains. LYSC-AVA39, encrypted in lysozyme C (UniProt ID: LYSC_HUMAN), displayed activity at 128 μmol L^−1^ against *P. aeruginosa* PA14 and both *E. coli* strains. Fragment CO7A1-AIG15 from human long-chain collage (UniProt ID: CO7A1_HUMAN) displayed activity at 32 μmol L^−1^ against *P. aeruginosa* PA14, while the protachykinin-1 (UniProt ID: TKN1_HUMAN) fragment TKN1-SSI27 was active at 64 μmol L^−1^ against *A. baumannii* ATCC19606. The physicochemical profiles of modern encrypted peptides (MEPs) are described in the Supplementary Discussion, Figs. 2 and S7, and Tables S7 and S8.

### Archaic encrypted peptides display antimicrobial activity in vitro

Six of 69 (8.7%) archaic protein fragments displayed *in vitro* antimicrobial activity (Fig. 3; Tables S10, S11). None of these fragments are reported as AMPs nor AMP subsequences in DBAASP (*23*). Fragment PDB6I34D-ALQ29 of chain D of the Neanderthal glycine decarboxylase protein displayed the broadest spectrum activity, moderately inhibiting both *P. aeruginosa* and *E. coli* strains (MICs from 32 to 128 μmol L^−1^). Denisovan transmembrane protein fragments A0A0S2IB02-AYT38 and A0A343EQH0-NVK38 displayed selective activity against *P. aeruginosa* PA01 at 128 μmol L^−1^. Similarly, A0A343AZS4-FMA25 encrypted within chain 1 of Denisovan NADH-ubiquinone oxidoreductase and A0A343EQH4-LAM11 from Neanderthal ATP synthase subunit A displayed selective activity against *A. baumannii* ATCC19606 at 128 μmol L^−1^. Neanderthal adenylosuccinate lyase fragment A0A384E0N4-DLI09 moderately inhibited *A. baumannii* ATCC19606 (128 μmol L^−1^), methicillin-resistant *Staphylococcus aureus* ATCC BAA-1556 (128 μmol L^−1^), and *Staphylococcus aureus* ATCC12600 (128 μmol L^−1^). The physicochemical profiles of the archaic encrypted peptides (AEP) are described in the Supplementary Discussion, Figs. 2 and S7, and Tables S8 and S12.

### Resistance to proteolytic degradation

Among MEPs, those curated for clustering strongly with known AMPs were highly resistant to serum proteases (Fig. 3). Up to 85% of the initial concentration of these peptides remained after six hours of continuous exposure to serum proteases. Shorter MEPs (8-residues long) were less susceptible to cleavage than longer MEPs (up to 24 residues), with ~35% of the initial concentration present after six hours of exposure to proteases versus 15–20%, respectively. On average, AEPs were more susceptible to proteolytic degradation than MEPs. An exception to this was the 9-residue-long encrypted peptide A0A384E0N4-DLI09, the shortest AEP tested. This short peptide resisted degradation for two hours, decreasing to 80% of its initial concentration, with ~55% of its initial concentration remaining after six hours of exposure (Fig. 3).

### Mechanism of action assays

MEPs and AEPs were investigated with fluorescent probes to determine how they affect the bacterial membrane. Positive control polymyxin B (PMB) is a peptide antibiotic having known permeabilizing and depolarizing effects (Figs. 3, S8, S9). In both assays, *A. baumannii* cells (Figs. 3, S8a-b, S8d-e) and *P. aeruginosa* PA01 (Fig. S8c,f) were exposed to the most active MEPs (CALR-GWT20, CBPZ-GSK4, TKN1-SSI27, and A7E2T1-SPR29 for *A. baumannii*) and AEPs (A0A384E0N4-DLI09 and A0A343EQH4-LAM11 for *A. baumannii;* A0A343EQH0-NVK38 and A0A0S2IB01-AYT38 for *P. aeruginosa*) at their respective MICs (Figs. 3, S8).

All MEPs except TKN1-SSI27 presented permeabilizing profiles similar to that of PMB. MEP TKN1-SSI27 initially demonstrated the slowest permeabilizing kinetics, yet progressively displayed the highest permeabilization efficiency (Figs. 3, S8, S9). The only peptide with an overall permeabilization efficacy lower than PMB was MEP CALR-GWT20. All MEPs initially displayed relatively slow depolarizing kinetics that increased over time. After 30 minutes, modern peptides had stronger depolarizing effects than PMB, which were maintained until the end of the experiment (Fig. 3). No significant differences were observed among their depolarizing activities.

AEPs permeabilized *A. baumannii* cells similarly to (A0A343EQH4-LAM11) or less than (A0A384E0N4-DLI09) PMB, but had much stronger depolarizing effects (Figs. 3, S8, S9). AEPs A0A343EQH0-NVK38 and A0A0S2IB01-AYT38 permeabilized *P. aeruginosa* cells (Figs. 3, S8), with higher relative fluorescence over time, indicating that *P. aeruginosa* was more sensitive than *A. baumannii* to these two peptides. Notably, A0A343EQH0-NVK38 and A0A0S2IB01-AYT38 were more strongly depolarizing than PMB for *P. aeruginosa* cells (Fig. S8).

### Anti-infective efficacy in preclinical animal models

To assess whether modern and archaic encrypted peptides retain their *in vitro* antimicrobial activity in complex living systems, we probed their antimicrobial properties in two mouse models (Fig. 1): a skin abscess model and a preclinical murine thigh infection model.

For skin abscess experiments, we selected MEPs and AEPs with activity at concentrations lower than 64 μmol L^−1^ against *A. baumannii* and *P. aeruginosa* PA01. Bacterial loads of 10^6^ and 10^5^ cells in 20 μL of *A. baumannii* and *P. aeruginosa* PA01, respectively, were administered to a skin abscess created on the back of each mouse. A single dose of PMB (control), MEP, or AEP was delivered as monotherapy to the infected area at MIC. Except for MEP A7E2T1-SPR39, all peptides demonstrated bactericidal effects in the skin abscess model (Fig. 4a). Activity levels were comparable to those of some of the most potent AMPs described to date in the literature using the same model, *i.e*., polybia-CP (*24*) and PaDBS1R6 (*25*). AEP A0A343EQH4-LAM11 and MEP CALR-GWT20 markedly reduced bacterial loads by 5–6 orders of magnitude against *A. baumannii*. AEPs A0A343EQH0-NVK38 and A0A0S2IB02-AYT38 reduced the bacterial load of *P. aeruginosa* by 3–4 orders of magnitude (Fig. 4b). No deleterious effects were observed in the animals (Fig. 4c).

**Fig. 4.**
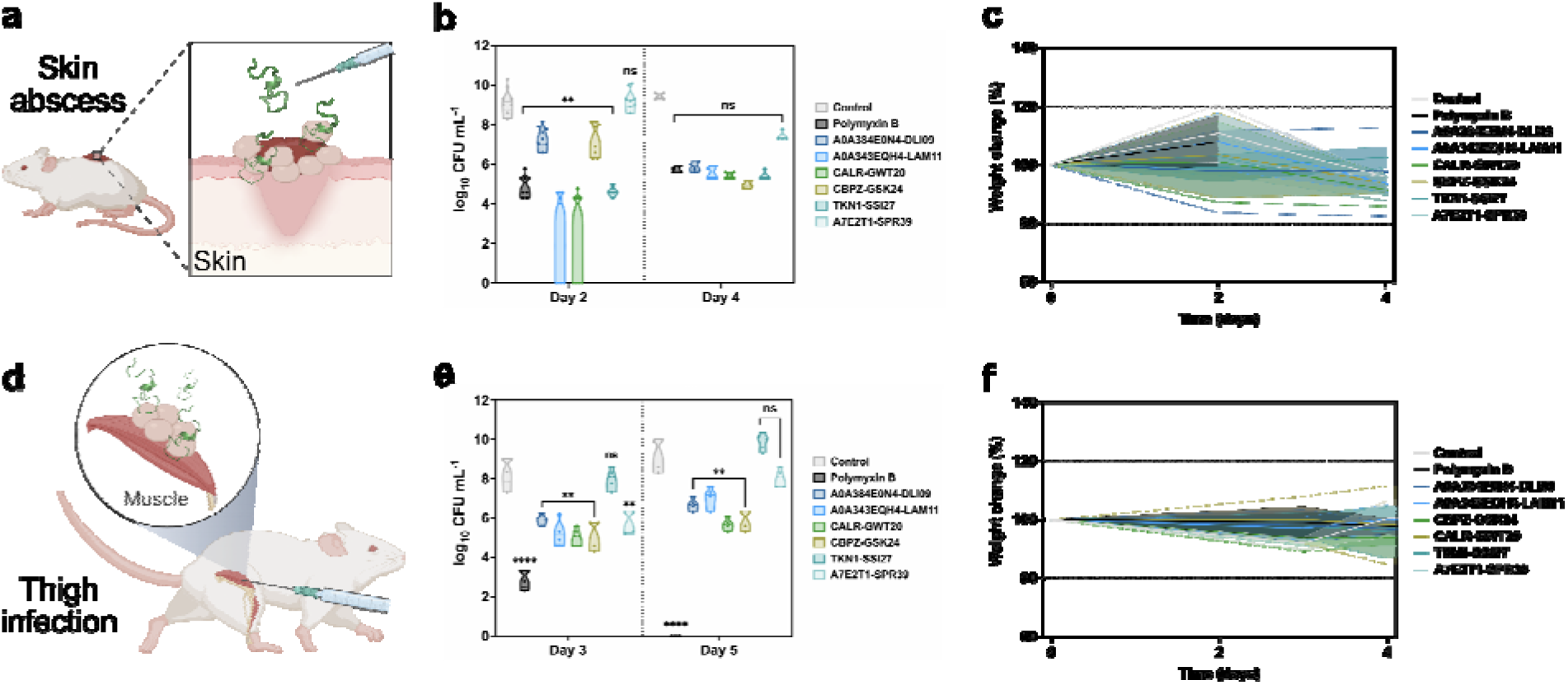
Anti-infective activity of modern and archaic encrypted peptides in pre-clinical animal models. **(a)** Schematic of the skin abscess mouse model used to assess the anti-infective activity of the modern and archaic encrypted peptides with activity against *A. baumannii* cells (***n*** = 8). **(b)** Peptides were tested at their MIC in a single dose one hour after the establishment of the infection. **(c)** To rule out toxic effects of the peptides, mouse weight was monitored throughout the whole extent of the experiment. **(d)** Schematic of the neutropenic thigh infection mouse model in which bacteria is injected intramuscularly in the right thigh and modern and archaic encrypted peptides were administered intraperitoneally to assess their systemic anti-infective activity (n□=□4). **(e)** All encrypted peptides, except TKN1-SSI17, showed bacteriostatic activity inhibiting proliferation of bacteria. Peptides with bacteriostatic activity were able to maintain their effect during the entire experiment (five days), except for A7E2T1-SPR39 that was effective for three days. **(f)** Mouse weight was monitored throughout the duration of the neutropenic thigh infection model (8 days total) to rule out potential toxic effects of cyclophosphamide injections, bacterial load, and the encrypted peptides. Statistical significance in **b** and **e** was determined using one-way ANOVA, **p□<□0.001, ****p□<D0.00001; features on the violin plots represent median and upper and lower quartiles. Data in **c** and **f** are the mean plus and minus the standard deviation. Figure created with BioRender.com and the PyMOL Molecular Graphics System, Version 2.1 Schrödinger, LLC.

For the preclinical murine thigh infection with *A. baumannii* (Fig. 4d), each peptide was injected at its MIC as a single intraperitoneal dose. The peptides used were active at concentrations lower than 64 μmol L^−1^ against *A. baumannii*. Three- and five-days post-treatment, all peptides tested presented bacteriostatic activity (Fig. 4e). In contrast, the PMB and levofloxacin controls displayed bactericidal activity and cleared the infection after five days. No significant changes in mouse weight were observed (Fig. 4f). As weight loss is a proxy for toxicity, these results suggest that the tested encrypted peptides are non-toxic.

## Discussion

This proof-of-concept study for ML-facilitated molecular de-extinction offers preliminary support for pharmacological prospection in paleoproteomes. We report the first known antimicrobial subsequences encrypted within archaic human proteins. While prior cleavage site classifiers favor protease-specific designs (*9–20*), the panCleave random forest is trained on protease-agnostic data yet is highly accurate for multiple specific proteases (Fig. 2). The panCleave pipeline uncovered six antimicrobial subsequences encrypted within extinct proteomes, allowing access to bioactive peptides with unusual amino acid distributions. Given the essential role of AMPs in innate immunity, host defense peptides derived from archaic introgression may have been retained in the modern human proteome. Potential maintenance of archaic AMPs in modern proteomes may merit future inquiry. The modern and archaic peptides presented here may offer new prototypes for antibiotic development.

The observed membrane depolarization was unexpected (*8*) and may have resulted from physicochemical differences between these peptides and human encrypted peptides mined with other computational methods (*8*), which do not depolarize bacterial cytoplasmic membranes. If encrypted peptides operate via mechanisms of action independent of cytoplasmic membrane depolarization, encrypted AMPs would be mechanistically distinct from non-encrypted AMPs. Encrypted AMP diversity is, therefore, an intriguing area for future inquiry.

### Rediscovery of a known antimicrobial motif

In addition to discovering new encrypted AMPs, the panCleave pipeline unintentionally uncovered a MEP containing a known bioactive subsequence. As lysozyme C is known to be bacteriolytic and to enhance immunoagent activity, it is unsurprising that a subsequence of this enzyme might itself display antimicrobial activity. A known encrypted peptide of human lysozyme C is a subsequence of panCleave-generated MEP LYSC-AVA39 (*26, 27*). The unintentional rediscovery of this antimicrobial motif in lysozyme C supports the use of the present pipeline for encrypted AMP discovery. Similarly, all encrypted AMPs discovered in the present work originate from proteins belonging to groups previously described in the encrypted peptide literature. As peptide fragments were not curated based on their precursor protein, this further lends support for panCleave as an encrypted AMP discovery tool.

### Precedents for modern precursor proteins

Secreted proteins have previously been targeted for bioactive encrypted peptide discovery (*28*). A prior whole-proteome search found an overrepresentation of secreted and membrane-bound proteins among encrypted AMP precursors, perhaps because AMPs are more likely to encounter pathogens in the extracellular environment (*8*). As has been thoroughly reviewed, enzymes are common precursors for encrypted host defense peptides (*29*). MEP precursor proteins identified in this study generally display catalytic activity (Table S9), with all MEP precursor groups having precedents in the encrypted peptide literature.

Proteases across the tree of life not only catalyze encrypted peptide release but also contain encrypted AMPs (*29*). In the present study, encrypted AMP CBPZ-GSK24 is derived from the protease carboxypeptidase Z, which may participate in prohormone processing (*30*).

Fragment CALR-GWT20 is derived from calreticulin, a calcium-binding chaperone protein that is highly conserved in multicellular life and is primarily localized to the endoplasmic reticulum. Calreticulin has been implicated in innate immune responses to bacterial infection in mammals, marine vertebrates, marine and terrestrial invertebrates, and plants (*31–34*). Vasostatin is a well-characterized anti-angiogenesis and anti-tumor encrypted peptide that is part of calreticulin (*35*), lending precedent for the presence of bioactive subsequences in this precursor.

Serine protease inhibitors in diverse marine organisms have displayed antibacterial and antiviral innate immunity functions (*36–38*). The observed antibacterial and antifungal activity of a kazal-type serine protease inhibitor in honeybee venom appears to act through microbial serine protease inhibition (*39*). MEP ISK5-GKI32 is encrypted within serine protease inhibitor kazal-type 5, which is known to yield encrypted peptides with protease inhibition activity when cleaved by the protease furin (*40*). The downregulation, deletion, and mutation of serine protease inhibitor kazal-type 5 are associated with inflammation, compromised skin-barrier function, atopic dermatitis, rosacea, and Netherton syndrome (*41–43*). Assaying ISK5-GKI32 against skin microbes implicated in these conditions could be a valuable area of future inquiry.

Oxidoreductases are known to contain encrypted AMPs in modern humans (*28, 29, 44*), *Bacillus* (*45*), *Desulfocurvibacter* (*46*), *Saccharomyces* (*47*), and *Physcomitrella* (*48*). In the present study, MEP XDH-AVA32 is a subsequence of the oxidoreductase xanthine dehydrogenase, which catalyzes the oxidative metabolism of purines. CO7A1-AIG15 is contained within the collagen alpha-1(VII) chain (syn. long-chain collagen), whose Gene Ontology molecular functions include serine-type endopeptidase inhibitor activity and extracellular matrix structural functionality (*49*).

MEP TKN1-SSI27 is contained within protachykinin-1, a neuropeptide implicated in antibacterial and antifungal humoral responses and defense responses to both Gram-negative and Gram-positive bacteria (*30*). A7E2T1-SPR29 originates from the uncharacterized protein fragment A7E2T1_HUMAN, which shares 99.21% identity with both the *Homo sapiens* neuropeptide W preproprotein (BLAST E-value 4e-78) and prepro-Neuropeptide W polypeptide (BLAST E-value 1e-77) (*50*). A7E2T1_HUMAN enables G protein-coupled receptor binding, according to Gene Ontology (*49*).

### Archaic precursors in the mitochondrial proteome

As publicly available Denisovan and Neanderthal data originate from the mitochondrial proteomes of these species, the AEP precursor proteins we identified are generally mitochondrial transmembrane proteins associated with transport, mitochondrial activity, and purine or ATP synthesis (Table S13). Precursor proteins were submitted to BLAST (*50*) to assess similarity to modern human analogs (Table S13). On average, the AEP precursor proteins shared 99.49% identity with a modern human protein (standard deviation < 0.003). All AEP precursors identified here belong to protein groups with precedents in the literature on encrypted host defense peptides, lending support for the use of panCleave for archaic human AMP prospection.

As discussed above, host defense peptides are known to be encrypted in oxidoreductases from across the kingdoms of life. AEP A0A343AZS4-FMA25 originated from the Denisovan transmembrane protein NADH-ubiquinone oxidoreductase chain 1 (EC 7.1.1.2), while A0A343EQH0-NVK38 is a subsequence of the 347-residue Neanderthal NADH-ubiquinone oxidoreductase chain 2 (EC 7.1.1.2). AEP A0A0S2IB02-AYT38 is a subsequence of the Denisovan cytochrome C oxidase subunit 1 (EC 7.1.1.9), a transmembrane protein that participates in the respiratory chain by catalyzing the reduction of oxygen to water.

Precedents for lyases as precursor proteins include an AMP enxcrypted within the pterin-4-alpha-carbinolamine dehydratase of *Mus musculus* (*46*). AEP A0A384E0N4-DLI09 is a subsequence of the Neanderthal adenylosuccinate lyase (syn. adenylosuccinase; EC 4.3.2.2), a coiled lyase involved in purine biosynthesis. AEP PDB6I34D-ALQ29 originates from chain D of the 984-residue Neanderthal lyase glycine decarboxylase.

The ATP synthase of the blowfly *Sarconesiopsis magellanica* is known to contain an encrypted AMP, and compounds excreted and secreted by this species have displayed antibacterial activity (*51*). Likewise, the Neanderthal ATP synthase subunit A was found to contain AEP A0A343EQH4-LAM11.

### Limitations of the study

The following limitations should be noted when interpreting the present work. The study design assumes that the extremely high similarity among the modern human, Neanderthal, and Denisovan proteomes is also suggestive of high conservation in protease activity (*e.g*., protease-substrate specificity). That is to say, we assume that a modern human protease with preference for a given amino acid sequence will also cleave Neanderthal or Denisovan proteins containing that subsequence. Though these assumptions leave claims of discovering naturally occurring archaic encrypted peptides unjustifiable, they do not undermine the present objective of bioinspired protein engineering. The construction of a synthetic negative dataset for training panCleave is also suboptimal, as negative sequences were not experimentally proven to be non-cleavage sites. In addition, the positive training data (*i.e*., observations that are cleavage sites) may be noisy, as the database from which they originate (*21*) is aggregated across diverse data sources.

## STAR Methods

- Key resources table
- Resource availability

- Lead contact
- Data and code availability
- Experimental model

- Bacterial strains and growth conditions
- Skin abscess infection mouse model
- Thigh infection mouse model
- Method details

- Antibacterial assays
- Membrane permeabilization assays
- Membrane depolarization assays
- Model training and testing data
- Hyperparameter tuning and model selection
- Modern protein fragment curation
- Archaic protein fragment curation

## Acknowledgments

Cesar de la Fuente-Nunez holds a Presidential Professorship at the University of Pennsylvania and acknowledges funding from the Procter & Gamble Company, United Therapeutics, a BBRF Young Investigator Grant, the Nemirovsky Prize, Penn Health-Tech Accelerator Award, and the Dean’s Innovation Fund from the Perelman School of Medicine at the University of Pennsylvania. Research reported in this publication was supported by the Langer Prize (AIChE Foundation), the National Institute of General Medical Sciences of the National Institutes of Health under award number R35GM138201 and the Defense Threat Reduction Agency (DTRA; HDTRA11810041 and HDTRA1-21-1-0014). Jacqueline R. M. A. Maasch acknowledges support from the University of Pennsylvania GAPSA-Provost Fellowship for Interdisciplinary Innovation and the Open Knowledge Foundation Frictionless Data for Reproducible Research Fellowship, funded by the Alfred P. Sloan Foundation. Computational resources included the Stampede2 supercomputer (Texas Advanced Computing Center, The University of Texas at Austin, TX, USA). We thank Dr. Mark Goulian for kindly donating the following strains: *Escherichia coli* AIC221 *[Escherichia coli* MG1655 phnE_2::FRT (control strain for AIC 222)] and *Escherichia coli* AIC222 *[Escherichia coli* MG1655 pmrA53 phnE_2::FRT (polymyxin resistant)]. We thank Dr. Karen Pepper for editing the manuscript and de la Fuente Lab members for insightful discussions. Figures created with BioRender.com are attributed as such. Molecules were rendered using the PyMOL Molecular Graphics System, Version 2.1 Schrödinger, LLC.

## Author contributions

Conceptualization: JRMAM, MDTT, MCRM, CFN

Methodology: JRMAM, MDTT, MCRM

Investigation: JRMAM, MDTT

Visualization: JRMAM, MDTT

Funding acquisition: CFN

Supervision: MCRM, CFN

Software: JRMAM

Formal analysis: JRMAM, MDTT

Writing – original draft: JRMAM, MDTT, CFN

Writing – review & editing: JRMAM, MDTT, MCRM, CFN

## STAR METHODS

### Key resources table

| Reagent or Resource | Source | Identifier |
| --- | --- | --- |
| **Chemicals** |  |  |
| Luria-Bertani broth | BD | 244620 |
| Tryptic soy broth | Sigma | T8907-1KG |
| Agar | Sigma | 05039 |
| MacConkey agar | RPI | M42560-500.0 |
| Phosphate buffer saline | Sigma | P3913-10PAK |
| Ammonium sulfate [(NH<sub>4</sub>)<sub>2</sub>SO<sub>4</sub>] | Chem Cruz | 7783-20-2 |
| Dipotassium hydrogen phosphate (K<sub>2</sub>HPO<sub>4</sub>) | Sigma | SLBR8555V |
| Monobasic potassium phosphate (KH<sub>2</sub>PO<sub>4</sub>) | Macron | 164500 |
| Iron (II) sulfate (FeSO<sub>4</sub>) | Amresco | 387 |
| Magnesium sulfate (MgSO<sub>4</sub>) | Amresco | 1333C215 |
| Glucose | Sigma | G5767 |
| 1-(N-phenylamino)naphthalene | Sigma | 104043 |
| 3,3'-dipropylthiadicarbocyanine iodide | Sigma | 43608 |
| HEPES | Fisher | BP310-100 |
| Potassium chloride (KCl) | Sigma | P3911 |
| **Software and Algorithms** |  |  |
| Python 3 | https://www.python.org/ |  |
| scikit-learn | https://scikit-learn.org/ |  |

### Resource availability

#### Lead contact

Further information and requests for resources should be directed to and will be fulfilled by the lead contact, Cesar de la Fuente-Nunez.

#### Data and code availability

All training data, testing data, and code used to develop the machine learning model are freely available on GitLab (https://gitlab.com/machine-biology-group-public/pancleave). All data pertaining to the experimental validation of generated peptides are available in the Supplementary Data.

### Experimental model

#### Bacterial strains and growth conditions

*Escherichia coli* ATCC11775, *Acinetobacter baumannii* ATCC19606, *Pseudomonas aeruginosa* PA01, *Pseudomonas aeruginosa* PA14, *Staphylococcus aureus* ATCC12600, *Staphylococcus aureus* ATCC BAA-1556 (methicillin-resistant strain), *Escherichia coli* AIC221 *[Escherichia coli* MG1655 phnE_2::FRT (control strain for AIC 222)] and *Escherichia coli* AIC222 *[Escherichia coli* MG1655 pmrA53 phnE_2::FRT (polymyxin resistant; colistin-resistant strain)], and *Klebsiella pneumoniae* ATCC13883 were grown and plated on Luria-Bertani (LB) or Pseudomonas Isolation (*Pseudomonas aeruginosa* strains) agar plates and incubated overnight at 37 °C from frozen stocks. After incubation, one isolated colony was transferred to 5 mL of medium (LB) or basal medium with glucose (BM2), and cultures were incubated overnight (16 h) at 37 °C. The following day, inocula were prepared by diluting the overnight cultures 1:100 in 5 mL of the respective media and incubating them at 37 °C until bacteria reached logarithmic phase (OD_600_ = 0.3 −0.5).

#### Skin abscess infection mouse model

*A. baumannii* ATCC19606 and *P. aeruginosa* PA01 were grown in tryptic soy broth (TSB) medium to an OD_600_ = 0.5. Next, cells were washed twice with sterile PBS (pH 7.4, 13,000 rpm for 1 min) and resuspended to a final concentration of 2×10^5^ and 5×10^6^ colony-forming units (CFU) mL^−1^ for *A. baumannii* and *P. aeruginosa*, respectively. Six-week-old female CD-1 mice were anesthetized with isoflurane for two minutes and had their backs shaved. A superficial linear skin abrasion was made with a needle to damage the stratum corneum and upper layer of the epidermis. An aliquot of 20 μL containing the bacterial load resuspended in PBS was inoculated over the scratched area. One hour after the infection, peptides diluted in water at their MIC value were administered to the infected area. Animals were euthanized and the area of scarified skin was excised two- and four-days post-infection, homogenized using a bead beater for 20 minutes (25 Hz), and 10-fold serially diluted for CFU quantification. Two independent experiments were performed with 4 mice per group in each condition.

#### Thigh infection mouse model

The mice were rendered neutropenic by two doses of cyclophosphamide (150 mg Kg^−1^) applied intraperitoneally with an interval of 72 h. One day after the last dose of cyclophosphamide, the mice were infected intramuscularly in their right thigh with a bacterial load of 10^6^ CFU mL^−1^ of *A. baumannii* ATCC19606. The bacteria were grown in tryptic soy broth (TSB), washed twice with PBS (pH 7.4), and resuspended to the desired concentration. Two hours later, peptides resuspended in water were administered intraperitoneally. Prior to each injection, mice were anesthetized with isoflurane and monitored for respiratory rate and pedal reflexes (*24, 52*). Next, we monitored the establishment of the infection and euthanized the mice. The infected area was excised two- and four-days post-infection, homogenized using a bead beater for 20 min (25 Hz), and 10-fold serially diluted for CFU quantification in MacConkey agar plates. The experiments were performed with 4 mice per group.

### Method details

#### Antibacterial assays

The 69 curated fragments were subjected to broth microdilution assays to assess *in vitro* antimicrobial activity. Minimum inhibitory concentration (MIC) values of the peptides were determined by using the broth microdilution technique with an initial inoculum of 5×10^6^ cells in LB or BM2 medium supplemented with glucose in nontreated polystyrene microtiter 96-well plates. Peptides were added to the plate as solutions in water at concentrations ranging from 0 to 128 μmol L^−1^. The MIC was considered as the lowest concentration of peptide that completely inhibited the visible growth of bacteria after 24 h of incubation of the plates at 37 °C. Plates were read in a spectrophotometer at 600 nm. All assays were done as three independent replicates.

#### Membrane permeabilization assays

The membrane permeability of the peptides was determined by using the 1-(N-phenylamino)naphthalene (NPN) uptake assay. NPN fluoresces weakly in extracellular environments and strongly when in contact with bacterial membrane lipids (Figs. 3e-g, S10a-c, and S11a), but only permeates the bacterial outer membrane when membrane integrity is compromised. *A. baumannii* ATCC19606 and *P. aeruginosa* PA01 were grown to an OD600 of 0.4, centrifuged (10,000 rpm at 4 °C for 10 min), washed, and resuspended in buffer (5 mmol L^−1^ HEPES, 5 mmol L^−1^ glucose, pH 7.4). Next, 4 μL of NPN solution (0.5 mmol L^−1^ – working concentration of 10 μmol L^−1^ after dilutions) was added to 100 μL of the bacterial solution in a white 96-well plate. The background fluorescence was recorded at λ_ex_ = 350 nm and λ_em_ = 420 nm. Peptide solutions in water (100 μL solution at their MIC values) were added to the 96-well plate, and fluorescence was recorded as a function of time until no further increase in fluorescence was observed (20 min).

#### Membrane depolarization assays

The ability of the peptides to depolarize the cytoplasmic membrane was determined by measurements of fluorescence of the membrane potential-sensitive dye, 3,3,-dipropyl thiadicarbocyanine iodide [DiSC_3_-(5)], a potentiometric fluorophore that fluoresces in response to an imbalance of the cytoplasmic membrane transmembrane potential (Fig. 3h-j, S10d-f, and S11b). Briefly, *A. baumannii* ATCC19606 and *P. aeruginosa* PA01 were grown at 37 °C with agitation until they reached mid-log phase (OD_600_ = 0.5). The cells were then centrifuged and washed twice with washing buffer (20 mmol L^−1^ glucose, 5 mmol L^−1^ HEPES, pH 7.2) and re-suspended to an OD_600_ of 0.05 in the same buffer containing 0.1 mol L^−1^ KCl. The cells (100 μL) were then incubated for 15 min with 20 nmol L^−1^ of DiSC_3_(5) until the reduction of fluorescence stabilized, indicating the incorporation of the dye into the bacterial membrane. Membrane depolarization was then monitored by observing the change in the fluorescence emission intensity of the membrane potential-sensitive dye, DiSC_3_-(5) (λ_ex_ = 622 nm, λ_em_ = 670 nm), after the addition of the peptides (100 μL solution at MIC values).

#### Model training and testing data

The panCleave random forest was trained and tested on all human protease substrates in the MEROPS Peptidase Database as of June 2020 (*21*). Substrate sequences for all human proteases available in MEROPS encompassed 369 proteases representing 6 catalytic types (Cysteine, Metallo, Serine, Aspartic, Threonine, and Mixed), 31 clans, and 73 families. Protease representation and amino acid frequency distributions for the MEROPS dataset are visualized in Figs. S2 and S3.

Model training and testing used a balanced dataset of 49,634 observations. Eight-residue cleavage site data were curated from the MEROPS Peptidase Database (*n* = 24,817 unique positive observations) (*21*) and combined with 8-residue sequences generated from the human proteome and random protein space (*n* = 24,817 unique negative observations). Redundant sequences, sites containing non-canonical amino acids, and sites of length shorter than 8 residues were removed from the positive dataset. Negative observations were generated by three methods, each constituting one third of the negative dataset: randomly selected 8-residue contiguous subsequences of the human proteome, randomly generated sequences adhering to the amino acid frequencies of the human proteome, and randomly generated sequences with no amino acid frequency constraints. No sequences were present in both the positive and negative datasets.

Training and 10-fold cross-validation were performed using 80% of total observations (*n* = 39,707). The remaining 20% of observations were reserved as an independent test set (*n* = 9,927). The train-test split was stratified by label to ensure that each split maintained a label distribution representative of the entire dataset. The complete training dataset, testing dataset, and Python code are available on GitLab and as supplemental files (https://gitlab.com/machine-biology-group-public/pancleave).

#### Hyperparameter tuning and model selection

Six classifiers were implemented using scikit-learn (https://scikit-learn.org/) and TensorFlow (https://www.tensorflow.org/): Gaussian Process (GP), K-Nearest Neighbor (KNN), Naive Bayes (NB), Random Forest (RF), Recurrent Neural Network (RNN), and Support Vector Machine (SVM). Each algorithm was trained and tested on 5 input representations: one-hot encoding, ProtFP (*53*), ST-Scale (*54*), Z-Scale (*55*), and UniRep (*56*). The resulting 30 candidate models each underwent Bayesian search hyperparameter tuning using the skopt Python package (https://scikit-optimize.github.io/) on the Stampede2 supercomputer (Texas Advanced Computing Center, The University of Texas at Austin, TX, USA).

Three tuned finalists were selected on the basis of superior test set accuracy: RF, RNN, and SVM, each trained on the ProtFP encoding. Finalists were assessed via three performance metrics, each computed using scikit-learn (https://scikit-learn.org/): test set accuracy, area under the receiver-operating characteristic curve (AUC-ROC), and average precision. Additionally, accuracy was assessed when thresholding the estimated probability of class membership at ≥50%, ≥60%, ≥70%, ≥80%, and ≥90%. The tradeoff between increases in accuracy and decreases in total valid observations at a given estimated probability threshold was quantified and visualized. Among the 30 candidate classifiers, an RF trained on the ProtFP protein encoding (*53*) was selected as the final model on the basis of marginally superior test set accuracy, AUC-ROC, average precision, and estimated probability thresholding.

#### Modern protein fragment curation

The panCleave pipeline was run on all modern human proteins tagged with the keyword “secreted” in UniProt (*30*) as of February 2021 (*n* = 3,676). Length distributions, amino acid frequencies, and PANTHER (http://www.pantherdb.org/) (*57*) classification data characterizing the modern secreted protein dataset are visualized (Figs. S2-S6). The initial 80,729 unique cleavage products were reduced to 3,738 fragments by filtering such that peptide lengths were between 8 and 40 residues, flanking cleavage sites were of an estimated probability of 0.8 or higher, and no fragments were subsequences of other fragments in the dataset.

Four curation methods were used to select panCleave-generated fragments for synthesis: 1) human expert curation; 2) ML model consensus using six publicly available AMP classifiers (*58–62*); 3) clustering against an in-house database of experimentally validated AMPs using CD-HIT-2D, an algorithm for sequence alignment and comparison of protein databases (*63*); and 4) random selection with no sampling bias. Twenty fragments were selected by each curation method (*n* = 80 total). In each case, fragment length was restricted to 8 to 40 amino acids.

Selection by a human expert entailed fully manual curation of 15 peptides predicted to be antimicrobial and 5 peptides predicted to be inactive. Consensus prediction used six publicly available ML-based AMP models: amPEPpy (https://github.com/tlawrence3/amPEPpy) (*58*), iAMPpred (http://cabgrid.res.in:8080/amppred/) (*59*), Macrel (https://www.big-data-biology.org/software/macrel/) (*60*), and three models available from AxPEP (https://app.cbbio.online/ampep/home) (*61, 62*). A positive consensus vote by at least three of these six models was required for selecting the 15 peptides predicted to be active. A negative consensus vote by all six models was required for selecting the 5 peptides predicted to be inactive. Random selection used no biasing criteria. The CD-HIT-2D clustering algorithm (http://weizhong-lab.ucsd.edu/cdhit-web-server/cgi-bin/index.cgi) (*63*) was used to rank fragments by percent similarity to an in-house dataset of experimentally validated AMPs, and the top 20 hits were selected as predicted AMPs for experimental validation.

#### Archaic protein fragment curation

The panCleave pipeline was run on all Neanderthal and Denisovan proteins available in UniProt (*30*) and NCBI (https://www.ncbi.nlm.nih.gov/protein/) as of February 2021 (*n* = 66 and *n* = 26, respectively). Six Neanderthal proteins (9.1%) and one Denisovan protein (3.8%) were identical to proteins in the modern proteome and were excluded. Results were filtered such that all fragments were between 8 and 40 residues in length and no fragments were subsequences of other fragments in the dataset. This filtering process yielded 249 unique Neanderthal cleavage products and 167 unique Denisovan cleavage products. No sequences were shared between modern human and Neanderthal panCleave results, nor between modern humans and Denisovans. There were 127 fragments common to both Neanderthals and Denisovans, leaving 289 non-redundant archaic fragments in total.

Archaic fragments were removed if present as subsequences of any protein in the modern human proteome. Archaic sequences were cross-referenced against all annotated and non-annotated modern human proteins (*n* = 75,552) and all isoforms (*n* = 40,403) available in UniProt as of February 2021. Subsequently, 73 archaic-only fragments remained (73/289, 25.3%). Of these, four were not selected for chemical synthesis because of their high hydrophobicity and aggregation propensity (*i.e*., WIGGQPVSYPFIIIG, VVAGVFLLIRFHPLA, LYDYGRWLVVVTGWTLFVGVYVVIE, and MTMYTTMTTLTLTSLIPPILTTLINPN), leaving 69 archaic-only fragments to be tested *in vitro*. All peptides used in the experiments were purchased from AAPPTec (Louisville, KY; USA).

