## Supplemental Information for "Molecular de-extinction of ancient antimicrobial peptides enabled by machine learning"

Jacqueline R. M. A. Maasch, Marcelo D. T. Torres, Marcelo C. R. Melo, Cesar de la Fuente-Nunez

**This PDF file includes:**

Supplementary discussion

Supplementary methods

Figs. S1 to S10

Tables S1 to S13

**Supplementary discussion:**

***A machine learning-based protein informatics pipeline for computational proteolysis***

Taking a bioinspired engineering approach to peptide discovery, we introduce a machine learning (ML) system that mines human proteomes for potential encrypted peptides. The panCleave Python pipeline is a protein informatics tool that uses ML to perform computational proteolysis: the *in silico* digestion of human proteins into peptide fragments (Figs. 1, S1). As the MEROPS database reports cleavage sites in terms of 8-residue P4:P4’ flanking sites (*1*), the panCleave random forest model predicts cleavage sites within each 8-residue subsequence of a given protein. Thus, the minimum length for an input protein is also 8 amino acids. The domain model for this Python pipeline is visualized in Fig. S1.

When presented with an 8-residue input, panCleave returns a binary label classifying the sequence as a cleavage site or non-cleavage site. Additionally, panCleave returns the estimated probability of class membership (i.e., the estimated probability that the assigned label for a given input is correct). Estimated probability reporting allows the user to filter predicted fragments by probability threshold, e.g., to bias fragment curation toward predictions with high estimated probability.

The panCleave pipeline performs the following procedure per input protein:

1. *Sliding window:* Every 8-residue contiguous subsequence of the protein sequence string is computed.
2. *Encoding:* Each subsequence is converted to a numerical feature vector using the ProtFP encoding method (*2*).
3. *Prediction:* The label and estimated probability of class membership are computed per subsequence.
4. *Fragmentation:* The full protein string is tokenized at each predicted cleavage site, yielding a list of peptide fragments.

Additional utility functions are also provided, including prediction filtering functionality and FASTA file conversion. Pipeline source code, tutorial notebooks, and documentation are available on GitLab (<https://gitlab.com/machine-biology-group-public/pancleave>).

***Modern encrypted peptide physicochemical profiles***

The modern encrypted peptides (MEPs) identified by panCleave were found to have similar content of aliphatic (Ala, Gly, Ile, Leu, Pro, and Val), aromatic (Phe, Trp, and Tyr), and basic (His, Arg, and Lys) amino acid residues as AMPs described in the DBAASP (*3*) (Fig. S7). Relative to encrypted peptides recently identified by a non-ML scoring function (*4*), MEPs identified here had 18% fewer basic residues and similar aliphatic and aromatic residue content, yet almost five-fold more acidic residues and 25% more polar residues (*4*). Interestingly, the MEPs displayed more acidic (Asp and Glu) and polar (Asn, Cys, Gln, Met, Ser, and Thr) amino acid residues than DBAASP AMPs (63% vs 26%, respectively). MEPs had notably higher median values for net charge and normalized hydrophobicity relative to archaic encrypted peptides (AEPs) (Fig. 2k,l), yet lower median values than recently reported encrypted AMPs (*4*) (Table S8). MEPs had slightly higher median length than AEPs and short AMPs (8–40 residues) reported in the DBAASP (*3*) (Fig. 2j), with MEPs being enriched in glycine and arginine relative to AEPs (Fig. 2i). The physicochemical properties of each individual MEP are described in Table S7.

***Archaic encrypted peptide physicochemical profiles***

The amino acid residue frequencies of AEPs differed from those of MEPs, recently described encrypted peptides (*4*), and known natural and synthetic AMPs described in the DBAASP (*3*) (Fig. 2). Similar to recently described encrypted peptides (*4*), AEPs presented 9% and 16% higher content of aromatic residues than MEPs and DBAASP AMPs, respectively. AEPs display 100% and 59% higher polar residue content than DBAASP AMPs and MEPs, respectively. AEPs presented the lowest frequency of basic residues across all groups analyzed, with frequencies 58%, 53%, and 62% lower than those of DBAASP AMPs, MEPs, and recently described encrypted AMPs. AEPs also contain many acidic residues, similar to MEPs. Aliphatic amino acid content was similar across all analyzed groups.

AEPs had lower median values for net charge and normalized hydrophobicity relative to MEPs, recently describe encrypted peptides (*4*), and AMPs reported in the DBAASP (*3*) (Figs. 2k,l; Tables S8, S13). AEPs were enriched in threonine, isoleucine, methionine, and phenylalanine (Fig. 2i) and demonstrated an extremely high median propensity to aggregate (Tables S8, S13). Although AEPs differed from MEPs and other encrypted peptides (Fig. S7), their properties were still within the range of some known AMPs due to their (i) high hydrophobic content, (ii) low net charge, (iii) disordered conformation, and (iv) considerable tendency to aggregate. Physicochemical properties of each individual AEP are described in Table S13.

***Positive hit rates by peptide curation method***

Protein fragment curation methods ranged in AMP hit rate, though none exceeded 20% accuracy. Activity was observed for 5% (1/20) of randomly selected peptides; 6.7% (1/15) of peptides predicted to be antimicrobial by a peptide chemistry expert; 20% (3/15) of peptides predicted to be antimicrobial by ML model consensus; and 15% (3/20) of peptides identified by CD-HIT-2D (*5*) (Fig. 2e). All ten sequences predicted to be inactive were indeed inactive against all pathogens tested experimentally (10/10, 100%). Small sample sizes aside, the marked difference in the accuracy of the curation method between predicted positives and predicted negatives might suggest that more is currently understood about non-antimicrobial protein space than about AMP diversity. Low agreement among popular, publicly available ML models for AMP discovery may suggest that models often overfit to their training-testing distributions, rather than being generalizable across antimicrobial protein space. Furthermore, both CD-HIT (*5*) and hierarchical *k*-means clustering (Fig. S10) failed to identify insightful boundaries between peptides with and without antimicrobial activity. In aggregate, these results emphasize the importance of characterizing antimicrobial protein space more extensively.

***Comparative performance evaluation for the machine learning model***

Direct comparative performance analyses were not possible for the panCleave model due to data availability, necessitating indirect comparisons (*i.e.,* comparing reported performance metrics rather than testing each model on the exact same evaluation dataset). As panCleave is the first known pan-protease cleavage classifier for human proteins, its overall accuracy cannot be benchmarked against an existing model. On the other hand, no existing protease-specific model was known to release exact training data, and very few models released test data. Training and testing data for prior models would be necessary to cross-reference against the panCleave data splits in order to prevent data leakage (*i.e.*, to ensure that testing instances were not present at training time), which could bias performance metrics in favor of a given model. The inability to directly compare panCleave with existing models highlights the importance of publicly releasing all training and testing data. Without the release of these data, the ML community is likely to face a significant reproducibility problem (*6*–*11*).

***Future questions for molecular de-extinction and evolutionary medicine***

Although the potential for molecular de-extinction to support drug discovery is a new open question, this proof-of-concept study for ML-facilitated molecular de-extinction offers preliminary support for the value of pharmacological prospection in paleoproteomes. Our results suggest that the de-extinction of archaic hominin peptides could expand the search space for protein therapies while remaining within the subspaces previously selected by human evolution. The potential for this approach to minimize pharmacological risks such as toxicity warrants investigation, as it may prove safer than mining evolutionarily distant or synthetic protein spaces.

The influence of archaic genomes on contemporary innate immunity could guide future interest in this line of inquiry. Encounters among modern humans, Neanderthals, and Denisovans resulted in multiple known admixture events that produced reproductively viable hybrid offspring (*12*, *13*). As Neanderthals and Denisovans evolved adaptive advantages in Eurasian environments over millennia, maintenance of introgressed alleles may have conferred fitness benefits to recently arrived modern humans (*14*). An overrepresentation of introgressed alleles has been observed in innate immunity genes relative to other genomic regions (*15*), resulting in a notable contribution to modern immune variation (*16*, *17*). Of current interest, the Neanderthal OAS1 haplotype has been seen to provide protection against COVID-19 infection in individuals of European descent (*18*), though an elevated risk of severe infection has also been linked to a 50 kb region of Neanderthal origin (*19*). Given their essential role in innate immunity, archaic host defense peptides may have entered the modern proteome. The present work does not pursue this question, though it may merit further inquiry. These questions of evolutionary medicine can be extended outside the *Homo* lineage, as has been done for ancient extant marsupial and monotreme host defense peptides (*20*).

**Supplementary Tables and Figures:**

***Supplementary Tables***

**Table S1. panCleave accuracy on proteases with at least 100 test set observations.** Total test set observations (*Total*) and total predictions with predicted labels that are concordant with true labels (*Concordant*) are noted. Mean probability refers to the predicted probability of class assignment averaged over all observations, as returned by the panCleave random forest (RF) model. Table is ordered by descending test set accuracy.

| **Protease MEROPS ID** | **Protease name** | **Concordant** | **Total** | **Mean probability** | **Accuracy** |
| --- | --- | --- | --- | --- | --- |
| C14.003 | Caspase-3 | 117 | 118 | 0.8266 | 0.9915 |
| C14.005 | Caspase-6 | 217 | 220 | 0.8635 | 0.9864 |
| S01.010 | Granzyme B | 300 | 322 | 0.7647 | 0.9317 |
| C13.004 | Legumain | 387 | 427 | 0.6714 | 0.9063 |
| C01.034 | Cathepsin S | 471 | 575 | 0.6403 | 0.8191 |
| C01.009 | Cathepsin V | 235 | 300 | 0.6362 | 0.7833 |
| C01.036 | Cathepsin K | 313 | 410 | 0.6353 | 0.7634 |
| M10.003 | Matrix metallopeptidase-2 | 353 | 471 | 0.6363 | 0.7495 |
| C01.032 | Cathepsin L | 406 | 543 | 0.6336 | 0.7477 |
| C01.060 | Cathepsin B | 75 | 101 | 0.6480 | 0.7426 |
| M10.005 | Matrix metallopeptidase-3 | 337 | 454 | 0.6264 | 0.7423 |
| A01.010 | Cathepsin E | 188 | 275 | 0.6101 | 0.6836 |
| A01.009 | Cathepsin D | 86 | 134 | 0.6180 | 0.6418 |
| S01.139 | Granzyme M | 86 | 136 | 0.6031 | 0.6324 |
| M12.004 | Meprin beta subunit | 108 | 183 | 0.6254 | 0.5902 |
| M12.002 | Meprin alpha subunit | 91 | 168 | 0.6232 | 0.5417 |

**Table S2. panCleave accuracy on protease families with at least 50 test set observations.** Protease family IDs are those reported in MEROPS (*1*). Total test set observations (*Total*) and total predictions with predicted labels that are concordant with true labels (*Concordant*) are noted. Mean probability refers to the predicted probability of class assignment averaged over all observations, as returned by the panCleave random forest (RF) model. Table is ordered by descending test set accuracy.

| **Protease family** | **Concordant** | **Total** | **Mean probability** | **Accuracy** |
| --- | --- | --- | --- | --- |
| C14 | 411 | 422 | 0.82512289 | 0.97393365 |
| C13 | 388 | 429 | 0.6712052 | 0.9044289 |
| S8 | 62 | 70 | 0.66046515 | 0.88571429 |
| C1 | 1026 | 1360 | 0.63477728 | 0.75441176 |
| M10 | 728 | 1008 | 0.63091031 | 0.72222222 |
| S1 | 616 | 872 | 0.67541674 | 0.70642202 |
| A1 | 270 | 433 | 0.61190775 | 0.62355658 |
| C2 | 56 | 98 | 0.63135532 | 0.57142857 |
| M12 | 179 | 328 | 0.62257116 | 0.54573171 |
| S26 | 34 | 69 | 0.59704888 | 0.49275362 |
| T1 | 17 | 51 | 0.64760432 | 0.33333333 |

**Table S3. panCleave accuracy on protease clans with at least 50 test set observations.** Protease clan IDs are those reported in MEROPS (*1*). Total test set observations (*Total*) and total predictions with predicted labels that are concordant with true labels (*Concordant*) are noted. Mean probability refers to the predicted probability of class assignment averaged over all observations, as returned by the panCleave random forest (RF) model. Table is ordered by descending test set accuracy.

| **Protease clan** | **Concordant** | **Total** | **Mean probability** | **Accuracy** |
| --- | --- | --- | --- | --- |
| CD | 785 | 836 | 0.7465014 | 0.93899522 |
| SB | 62 | 70 | 0.66046515 | 0.88571429 |
| CA | 1081 | 1459 | 0.63442325 | 0.74091844 |
| PA | 616 | 872 | 0.67541674 | 0.70642202 |
| MA | 910 | 1346 | 0.62844619 | 0.67607727 |
| AA | 270 | 433 | 0.61190775 | 0.62355658 |
| SF | 34 | 69 | 0.59704888 | 0.49275362 |
| PB | 18 | 52 | 0.65295276 | 0.34615385 |

**Table S4. Protease-specific accuracy of panCleave as compared to published cleavage site models.** The following table is adapted from Table S6 in (*21*). Reported panCleave values are test set accuracy. Best accuracy per protease is represented in bold type. Note that all comparisons are relative, with accuracies as reported in (*21*); direct comparisons were not possible, as training and testing data were not released for prior models. Models are Cascleave (*22*), CAT3 (*23*), CleavPredict (*24*), ScreenCap3 (*25*), SitePrediction (*26*), PROSPERous (*27*), and DeepCleave (*21*). Results for DeepCleave are reported with and without transfer learning (TL), as reported in the original publication.

| **Protease name** | **MEROPS ID** | **panCleave** | **Cascleave** | **CAT3** | **CleavPredict** | **ScreenCap3** | **SitePrediction** | **PROSPERous** | **DeepCleave (without TL)** | **DeepCleave (with TL)** |
| --- | --- | --- | --- | --- | --- | --- | --- | --- | --- | --- |
| Caspase-1 | C14.001 | **0.9268** | 0.5119 | 0.6786 | - | 0.7368 | 0.8571 | 0.8750 | 0.8315 | 0.9205 |
| Caspase-3 | C14.003 | **0.9915** | 0.5920 | 0.7731 | - | 0.8741 | 0.9275 | 0.9656 | 0.9759 | 0.9856 |
| Caspase-7 | C14.004 | 0.9524 | 0.6143 | 0.8378 | - | 0.8684 | 0.9605 | 0.9359 | 0.9625 | **0.9744** |
| Caspase-6 | C14.005 | 0.9864 | 0.5286 | 0.6583 | - | 0.7802 | 0.8767 | 0.9850 | **0.9935** | 0.9901 |
| Caspase-2 | C14.006 | **1.0000** | 0.5519 | 0.7632 | - | 0.8333 | - | - | 0.9878 | 0.9880 |
| Matrix metallopeptidase-8 | M10.002 | 0.6667 | - | - | 0.7273 | - | **0.8750** | 0.8250 | 0.6250 | 0.7879 |
| Matrix metallopeptidase-2 | M10.003 | 0.7495 | - | - | 0.6032 | - | 0.7133 | 0.8736 | - | **0.8899** |
| Matrix metallopeptidase-9 | M10.004 | 0.6613 | - | - | 0.6081 | - | 0.8378 | 0.8077 | 0.5734 | **0.8613** |
| Matrix metallopeptidase-3 | M10.005 | 0.7423 | - | - | 0.7273 | - | 0.8571 | **0.8810** | 0.7027 | 0.8780 |
| Matrix metallopeptidase-7 | M10.008 | 0.7805 | - | - | - | - | 0.7375 | 0.8523 | 0.6264 | **0.9318** |
| Matrix metallopeptidase-12 | M10.009 | 0.7143 | - | - | - | - | 0.6786 | 0.8378 | 0.6316 | **0.9079** |
| Matrix metallopeptidase-1 | M10.014 | 0.4815 | - | - | - | - | - | 0.8125 | 0.6275 | **0.8519** |

**Table S5. Modern secreted protein fragments with antimicrobial activity in BM2 medium with glucose.** Minimum inhibitory concentration (MIC) values (μmol L^-1^) for peptides screened against 4 pathogenic strains (dash indicates no activity). If no peptides demonstrated activity against a tested strain, the strain is not included here. Curation methods are machine learning model consensus vote (ML), random selection (RS), and human expert (HE). Predicted label indicates predicted antimicrobial activity (1), no predicted activity (0), or no prediction (NA).

| **Fragment ID** | **Fragment sequence** | **Length**  **(number of amino acid residues)** | **Curation method** | **Predicted label** | ***P. aeruginosa* PA01**  MIC (μmol L^-1^) | ***P. aeruginosa* PA14**  MIC (μmol L^-1^) | ***E. coli* AIC221**  MIC (μmol L^-1^) | ***E. coli* AIC222**  MIC (μmol L^-1^) |
| --- | --- | --- | --- | --- | --- | --- | --- | --- |
| CBPZ-GSK24 | GSKPWWWSYFTSLSTHRPRWLLKY | 24 | ML | 1 | 8 | 4 | 4 | 2 |
| XDH-AVA32 | AVAKLPAQKTEVFRGVLEQLRWFA  GKQVKSVA | 32 | ML | 1 | - | - | 32 | 32 |
| LYSC-AVA39 | AVACAKRVVRDPQGIRAWVAWRN  RCQNRDVRQYVQGCGV | 39 | ML | 1 | - | 128 | 128 | 128 |
| ISK5-GKI32 | GKIHGNTCSMCEAFFQQEAKEKERAEPRAKVK | 32 | RS | NA | - | - | 128 | 128 |
| CALR-GWT20 | GWTSRWIESKHKSDFGKFVL | 20 | HE | 1 | - | - | - | 128 |

**Table S6. Modern secreted protein fragments with antimicrobial activity in LB medium.** Minimum inhibitory concentration (MIC) values (μmol L^-1^) for peptides screened against 4 pathogenic strains (dash indicates no activity). If no peptides demonstrated activity against a tested strain, the strain is not reported here. Curation methods are machine learning model consensus vote (ML), random selection (RS), human expert (HE), and CD-HIT (CD). All modern secreted protein fragments were predicted as antimicrobials (Predicted label = 1).

| **Fragment ID** | **Fragment sequence** | **Fragment**  **length** | **Curation method** | **Predicted label** | ***A. baumannii* ATCC19606**  MIC (μmol L^-1^) | ***P. aeruginosa PA14***  MIC (μmol L^-1^) | ***E. coli* AIC221**  MIC (μmol L^-1^) | ***E. coli* AIC222**  MIC (μmol L^-1^) |
| --- | --- | --- | --- | --- | --- | --- | --- | --- |
| CBPZ-GSK24 | GSKPWWWSYFTSLSTHRPRWLLKY | 24 | ML | 1 | 16 | - | - | - |
| CALR-GWT20 | GWTSRWIESKHKSDFGKFVL | 20 | HE | 1 | 64 | - | - | - |
| CO7A1-AIG15 | AIGPKGDRGFPGPLG | 15 | CD | 1 | - | 32 | - | - |
| TKN1-SSI27 | SSIEKQVALLKALYGHGQISHKRHKTD | 27 | CD | 1 | 64 | - | - | - |
| A7E2T1-SPR29 | SPRYHTVGRAAGLLMGLRRSPYLWRRALR | 29 | CD | 1 | 8 | - | 64 | 64 |

**Table S7. Properties of modern encrypted peptides.** Medians are reported with standard deviations in parentheses. Physicochemical properties were calculated using the DBAASP (*3*) (<https://dbaasp.org/tools?page=property-calculation>) and the Eisenberg and Weiss hydrophobicity scale (*28*). Reported properties are: Normalized Hydrophobic Moment (NHM), Normalized Hydrophobicity (NH), Net Charge (NC), Isoelectric Point (IP), Penetration Depth (PD), Tilt Angle (TA), Disordered Conformation Propensity (DCP), Linear Moment (LM), Propensity to *in vitro* Aggregation (PA), Angle Subtended by the Hydrophobic Residues (AS), Amphiphilicity Index (AI), and Propensity to PPII coil (PC).

| **Fragment ID** | **NHM** | **NH** | **NC** | **IP** | **PD** | **TA** | **DCP** | **LM** | **PA** | **AS** | **AI** | **PC** |
| --- | --- | --- | --- | --- | --- | --- | --- | --- | --- | --- | --- | --- |
| **CBPZ-GSK24** | 0.11 | 0.02 | 4 | 10.87 | 15 | 90 | -0.19 | 0.31 | 356.51 | 40 | 2.15 | 1.13 |
| **XDH-AVA32** | 0.32 | -0.02 | 4 | 11.07 | 18 | 114 | 0.12 | 0.14 | 0.00 | 90 | 1.03 | 0.95 |
| **LYSC-AVA39** | 0.28 | 0.25 | 6 | 10.98 | 30 | 141 | -0.06 | 0.24 | 12.06 | 60 | 1.15 | 1.01 |
| **ISK5-GKI32** | 0.18 | 0.30 | 2 | 8.93 | 30 | 42 | -0.16 | 0.31 | 0.00 | 30 | 1.05 | 0.97 |
| **CALR-GWT20** | 0.10 | 0.06 | 2 | 10.43 | 24 | 159 | -0.08 | 0.33 | 0.00 | 40 | 1.50 | 0.96 |
| **CO7A1-AIG15** | 0.33 | -0.14 | 1 | 10.17 | 22 | 140 | 0.06 | 0.50 | 0.00 | 170 | 0.41 | 1.01 |
| **TKN1-SSI27** | 0.14 | 0.15 | 3 | 10.39 | 19 | 70 | -0.11 | 0.33 | 4.11 | 40 | 1.12 | 0.95 |
| **A7E2T1-SPR29** | 0.27 | 0.23 | 7 | 12.23 | 16 | 97 | -0.15 | 0.31 | 1.58 | 50 | 1.23 | 1.02 |
| **Median**  **(SD)** | **0.23**  **(0.09)** | **0.11**  **(0.15)** | **3.50 (2.07)** | **10.65 (0.94)** | **20.50 (5.87)** | **105.50 (39.6)** | **-0.10 (0.11)** | **0.31**  **(0.1)** | **0.79 (125.22)** | **45**  **(46.29)** | **1.14**  **(0.49)** | **0.99 (0.06)** |

**Table S8. Physicochemical properties of archaic, modern, and recently described encrypted peptides** (*4*) **and AMPs listed in the DBAASP** (*3*)**.** Medians are reported with standard deviations in parentheses. Physicochemical properties were calculated using the DBAASP (<https://dbaasp.org/tools?page=property-calculation>) and the Eisenberg and Weiss hydrophobicity scale (*28*). Reported properties are: Normalized Hydrophobic Moment (NHM), Normalized Hydrophobicity (NH), Net Charge (NC), Isoelectric Point (IP), Penetration Depth (PD), Tilt Angle (TA), Disordered Conformation Propensity (DCP), Linear Moment (LM), Propensity to *in vitro* Aggregation (PA), Angle Subtended by the Hydrophobic Residues (AS), Amphiphilicity Index (AI), and Propensity to PPII coil (PC).

| **Antimicrobial peptides** | **NHM** | **NH** | **NC** | **IP** | **PD** | **TA** | **DCP** | **LM** | **PA** | **AS** | **AI** | **PC** |
| --- | --- | --- | --- | --- | --- | --- | --- | --- | --- | --- | --- | --- |
| **Archaic encrypted AMPs**  *(n = 6)* | 0.10  (0.18) | -0.35 (0.38) | 1  (2.53) | 6.93 (3.86) | 13  (9.67) | 77  (45.36) | 0.41  (0.3) | 0.33 (0.15) | 550.22 (612.11) | 75 (119.94) | 0.61 (0.19) | 0.99 (0.07) |
| **Modern encrypted AMPs**  *(n = 8)* | 0.23  (0.09) | 0.11  (0.15) | 3.50 (2.07) | 10.65 (0.94) | 20.50 (5.87) | 105.50 (39.6) | -0.10 (0.11) | 0.31  (0.1) | 0.79 (125.22) | 45  (46.29) | 1.14  (0.49) | 0.99 (0.06) |
| **Previously reported encrypted AMPs** (*4*)  *(n = 35)* | 0.23  (0.15) | 0.20  (0.19) | 7  (3.34) | 11.58  (0.78) | 21  (7.19) | 88  (30.89) | -0.17  (0.21) | 0.23  (0.05) | 0  (127.27) | 60  (31.48) | 1.23  (0.31) | 1.03  (0.12) |
| **AMPs in the DBAASP** (*3*)  *(n = 14,995)* | 0.37  (0.24) | 0.07  (0.42) | 4  (3.18) | 11.15  (2.21) | 15  (6.98) | 88  (33.21) | -0.05  (0.46) | 0.29  (0.11) | 0  (149.89) | 110  (69.95) | 1.23  (0.91) | 0.97  (0.13) |

**Table S9. Precursor proteins of modern human panCleave fragments with antimicrobial activity.** Precursor protein data obtained from the PANTHER Classification System (<http://www.pantherdb.org/>) (*7*). Dashes indicate that data were unavailable.

| **ID** | **Fragment sequence** | **Gene ID** | **Precursor**  **UniProt ID** | **Precursor gene name** | **PANTHER Protein Class** |
| --- | --- | --- | --- | --- | --- |
| CALR-GWT20 | GWTSRWIESKHKSDFGKFVL | HUMAN\|HGNC=1455\|UniProtKB=P27797 | CALR_HUMAN | Calreticulin | chaperone |
| XDH-AVA32 | AVAKLPAQKTEVFRGVLE  QLRWFAGKQVKSVA | HUMAN\|HGNC=12805\|UniProtKB=P47989 | XDH_HUMAN | Xanthine dehydrogenase/oxidase | oxidoreductase |
| ISK5-GKI32 | GKIHGNTCSMCEAFFQQE  AKEKERAEPRAKVK | HUMAN\|HGNC=15464\|UniProtKB=Q9NQ38 | ISK5_HUMAN | Serine protease inhibitor Kazal-type 5 | protease inhibitor |
| CBPZ-GSK24 | GSKPWWWSYFTSLST  HRPRWLLKY | HUMAN\|HGNC=2333\|UniProtKB=Q66K79 | CBPZ_HUMAN | Carboxypeptidase Z | protease |
| LYSC-AVA39 | AVACAKRVVRDPQGIRA  WVAWRNRCQNRDVRQY  VQGCGV | HUMAN\|HGNC=6740\|UniProtKB=P61626 | LYSC_HUMAN | Lysozyme C | - |
| CO7A1-AIG15 | AIGPKGDRGFPGPLG | HUMAN\|HGNC=2214\|UniProtKB=Q02388 | CO7A1_HUMAN | Collagen alpha-1(VII) chain (Long-chain collagen) (LC collagen) | extracellular matrix structural protein |
| TKN1-SSI27 | SSIEKQVALLKALYGHGQIS  HKRHKTD | HUMAN\|HGNC=11517\|UniProtKB=P20366 | TKN1_HUMAN | Protachykinin-1 (PPT) [Cleaved into: Substance P; Neurokinin A (NKA) (Neuromedin L) (Substance K); Neuropeptide K (NPK); Neuropeptide gamma; C-terminal-flanking peptide] | - |
| A7E2T1-SPR29 | SPRYHTVGRAAGLLMGLRR  SPYLWRRALR | - | A7E2T1_HUMAN | Uncharacterized protein (Fragment) | - |

**Table S10. Archaic encrypted peptides with antimicrobial activity in BM2 medium.** Minimum inhibitory concentration (MIC) values (μmol L^-1^) for 4 pathogenic strains. If no peptides demonstrated activity against a tested strain, the strain is not reported here.

| **Fragment ID** | **Fragment sequence** |  | **MIC (μmol L^-1^)** | | | |
| --- | --- | --- | --- | --- | --- | --- |
|  |  | **Length** | ***P. aeruginosa* PA01** | ***P. aeruginosa* PA14** | ***E. coli* AIC221** | ***E. coli* AIC222** |
| PDB6I34D-ALQ29 | ALQLCYRHNKRRKFFVDPRCHPQTIAVVQ | 29 | 64 | 32 | 128 | 128 |

| **Fragment ID** | **Fragment sequence** | **Length** | ***P. aeruginosa* PA01** | ***S. aureus* ATCC12600** | ***A. baumannii* ATCC19606** | ***S. aureus* ATCC BAA-1556 - MRSA** |
| --- | --- | --- | --- | --- | --- | --- |
| A0A384E0N4-DLI09 | DLIERIQAD | 9 | - | 128 | 128 | 128 |
| A0A343EQH4-LAM11 | LAMVIPLWAGA | 11 | - | - | 128 | - |
| A0A343AZS4-FMA25 | FMAEYTNIIMMNTLTTTIFLGTTYN | 25 | - | - | 128 | - |
| A0A343EQH0-NVK38 | NVKMKWQFEHTKPTPFLPTLITLTTLLLPISPFMLMIL | 38 | 128 | - | - | - |
| A0A0S2IB02-AYT38 | AYTTWNILSSAGSFISLTAVMLMIFMIWEAFASKRKVL | 38 | 128 | - | - | - |

**Table S11. Archaic encrypted peptides with antimicrobial activity in LB medium.** Minimum inhibitory concentration (MIC) values (μmol L^-1^) for 4 pathogenic strains (dash indicates no activity). If no peptides demonstrated activity against a tested strain, the strain is not reported here.

**Table S12. Properties of archaic encrypted peptides.** Medians are reported with standard deviations in parentheses. Physicochemical properties were calculated using the DBAASP (*3*) (<https://dbaasp.org/tools?page=property-calculation>) and the Eisenberg and Weiss hydrophobicity scale (*28*). Reported properties are: Normalized Hydrophobic Moment (NHM), Normalized Hydrophobicity (NH), Net Charge (NC), Isoelectric Point (IP), Penetration Depth (PD), Tilt Angle (TA), Disordered Conformation Propensity (DCP), Linear Moment (LM), Propensity to *in vitro* Aggregation (PA), Angle Subtended by the Hydrophobic Residues (AS), Amphiphilicity Index (AI), and Propensity to PPII coil (PC).

| **Fragment ID** | **NHM** | **NH** | **NC** | **IP** | **PD** | **TA** | **DCP** | **LM** | **PA** | **AS** | **AI** | **PC** |
| --- | --- | --- | --- | --- | --- | --- | --- | --- | --- | --- | --- | --- |
| PDB6I34D-ALQ29 | 0.07 | 0.23 | 5 | 10.75 | 30 | 16 | -0.1 | 0.34 | 208.02 | 50 | 0.99 | 1.09 |
| A0A384E0N4-DLI09 | 0.54 | 0.16 | -2 | 3.57 | 17 | 72 | 0.33 | 0.31 | 0.00 | 130 | 0.55 | 0.93 |
| A0A343EQH4-LAM11 | 0.18 | -0.77 | 0 | 3.5 | 5 | 82 | 0.82 | 0.00 | 0.00 | 360 | 0.63 | 0.99 |
| A0A343AZS4-FMA25 | 0.06 | -0.35 | -1 | 3.22 | 13 | 89 | 0.43 | 0.27 | 1043.42 | 60 | 0.46 | 0.98 |
| A0A343EQH0-NVK38 | 0.10 | -0.34 | 2 | 10.38 | 13 | 148 | 0.38 | 0.42 | 892.42 | 50 | 0.58 | 1.11 |
| A0A0S2IB02-AYT38 | 0.09 | -0.39 | 2 | 10.28 | 3 | 40 | 0.48 | 0.39 | 1447.14 | 90 | 0.79 | 0.99 |
| **Median**  **(SD)** | **0.10**  **(0.18)** | **-0.35 (0.38)** | **1**  **(2.53)** | **6.93 (3.86)** | **13**  **(9.67)** | **77**  **(45.36)** | **0.41**  **(0.3)** | **0.33 (0.15)** | **550.22 (612.11)** | **75 (119.94)** | **0.61 (0.19)** | **0.99 (0.07)** |

**Table S13. Precursor proteins of archaic encrypted peptides with antimicrobial activity.** Precursor protein data obtained from UniProt (*29*) and NCBI Protein Databases (<https://www.ncbi.nlm.nih.gov/protein/>). Percent identity shared with a modern human protein (% ID) and query coverage (QC) were computed by BLAST analysis (*30*), with values reported for the top modern human hits.

| **Precursor name** | **Hominin** | **UniProt ID** | **NCBI ID** | **% ID**  **(QC)** | **Length** | **Keywords** | **Features** | **Fragment ID** | **Fragment sequence** |
| --- | --- | --- | --- | --- | --- | --- | --- | --- | --- |
| Chain D, Neanderthal Glycine decarboxylase | *Homo sapiens neanderthalensis* (Neanderthal) |  | gi\|1777435468\|pdb\|6I34\|D | 99.90%  (100%) | 984 |  |  | PDB6I34D-ALQ29 | ALQLCYRHNKRRKFFV  DPRCHPQTIAVVQ |
| Adenylosuccinate lyase (ASL) (EC 4.3.2.2) (Adenylosuccinase) | *Homo sapiens neanderthalensis* (Neanderthal) | A0A384E0N4 | gi\|1393955578\|pdb\|5NXA\|H | 99.59%  (100%) | 487 | 3D-structure;  Coiled coil;  Lyase;  Purine biosynthesis | Coiled coil (1); Domain (1) | A0A384E0N4-DLI09 | DLIERIQAD |
| ATP synthase subunit a | *Homo sapiens neanderthalensis* (Neanderthal) | A0A343EQH4 | gi\|1214786277\|gb\|ASK06270.1\| | 99.56%  (100%) | 226 | ATP synthesis;  CF(0);  Hydrogen ion transport;  Ion transport;  Membrane;  Mitochondrion;  Mitochondrion inner membrane;  Transmembrane;  Transmembrane helix;  Transport | Transmembrane (6) | A0A343EQH4-LAM11 | LAMVIPLWAGA |
| NADH-ubiquinone oxidoreductase chain 1 (EC 7.1.1.2) | *Homo sapiens* subsp. 'Denisova' (Denisova hominin) | A0A343AZS4 | gi\|1141958635\|gb\|AQD17584.1\| | 99.06%  (100%) | 318 | Membrane;  Mitochondrion;  NAD;  Transmembrane;  Transmembrane helix;  Transport;  Ubiquinone | Transmembrane (8) | A0A343AZS4-FMA25 | FMAEYTNIIMMNTLTTT  IFLGTTYN |
| NADH-ubiquinone oxidoreductase chain 2 (EC 7.1.1.2) | *Homo sapiens neanderthalensis* (Neanderthal) | A0A343EQH0 | gi\|1578894740\|gb\|ASK06266.2\| | 99.38% (92%) | 347 | Electron transport;  Membrane;  Mitochondrion;  Mitochondrion inner membrane;  NAD;  Respiratory chain;  Translocase;  Transmembrane;  Transmembrane helix;  Transport;  Ubiquinone | Domain (2); Transmembrane (8) | A0A343EQH0-NVK38 | NVKMKWQFEHTKPTP  FLPTLITLTTLLLPISPF  MLMIL |
| Cytochrome c oxidase subunit 1 (EC 7.1.1.9) | *Homo sapiens* subsp. 'Denisova' (Denisova hominin) | A0A0S2IB02 | gi\|1141958637\|gb\|AQD17586.1\| | 99.42%  (100%) | 513 | Calcium;  Copper;  Electron transport;  Heme;  Iron;  Magnesium;  Membrane;  Metal-binding;  Mitochondrion;  Mitochondrion inner membrane;  Respiratory chain;  Sodium;  Translocase;  Transmembrane;  Transmembrane helix;  Transport | Domain (1); Transmembrane (12) | A0A0S2IB02-AYT38 | AYTTWNILSSAGSFIS  LTAVMLMIFMIWEAF  ASKRKVL |

***Supplementary Figures***

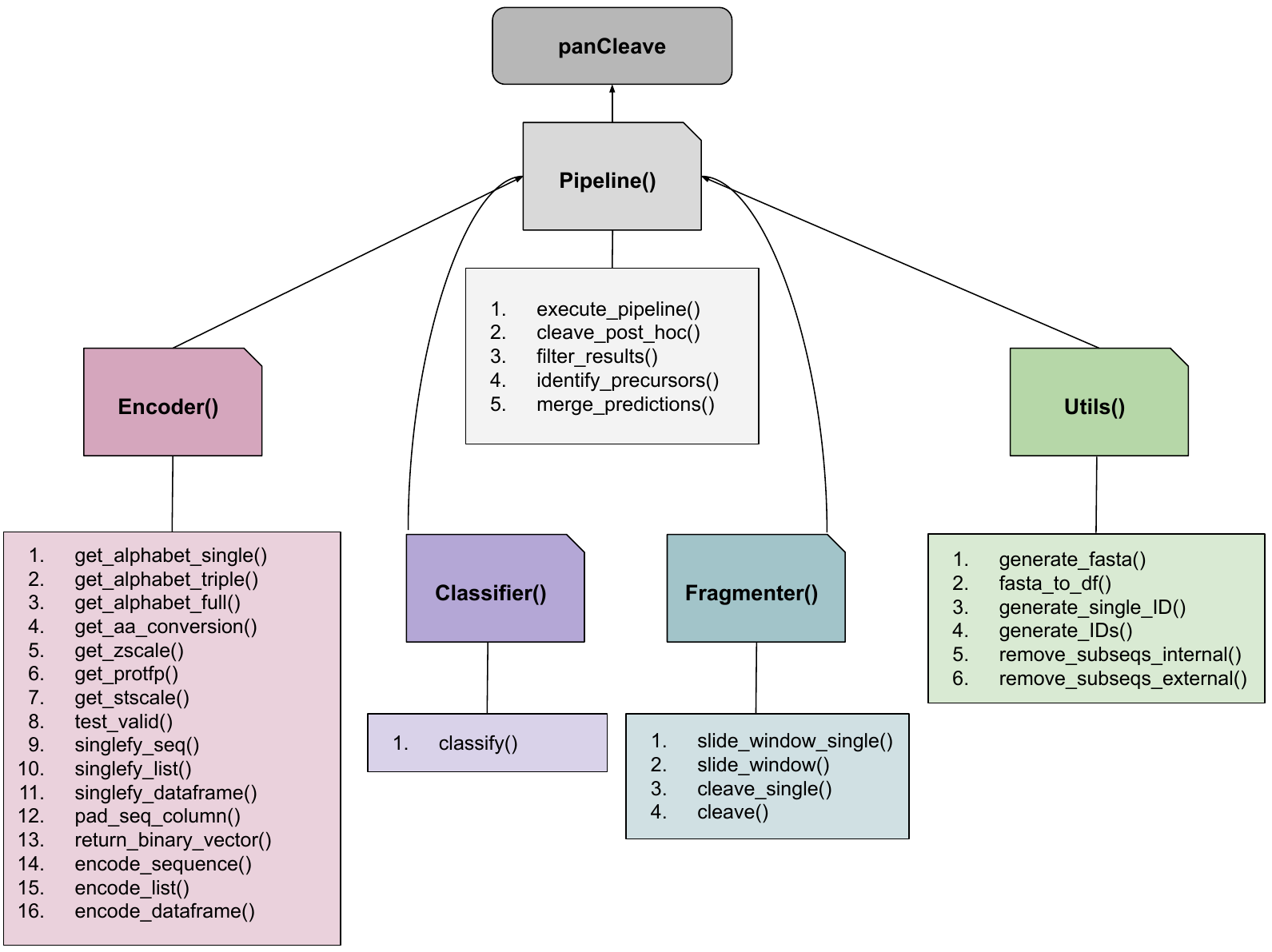

**Fig. S1. Domain model for the panCleave pipeline in Python.** The class *Pipeline* is dependent on classes *Encoder*, *Classifier*, *Fragmenter*, and *Utils*. Each class features the methods enumerated here.

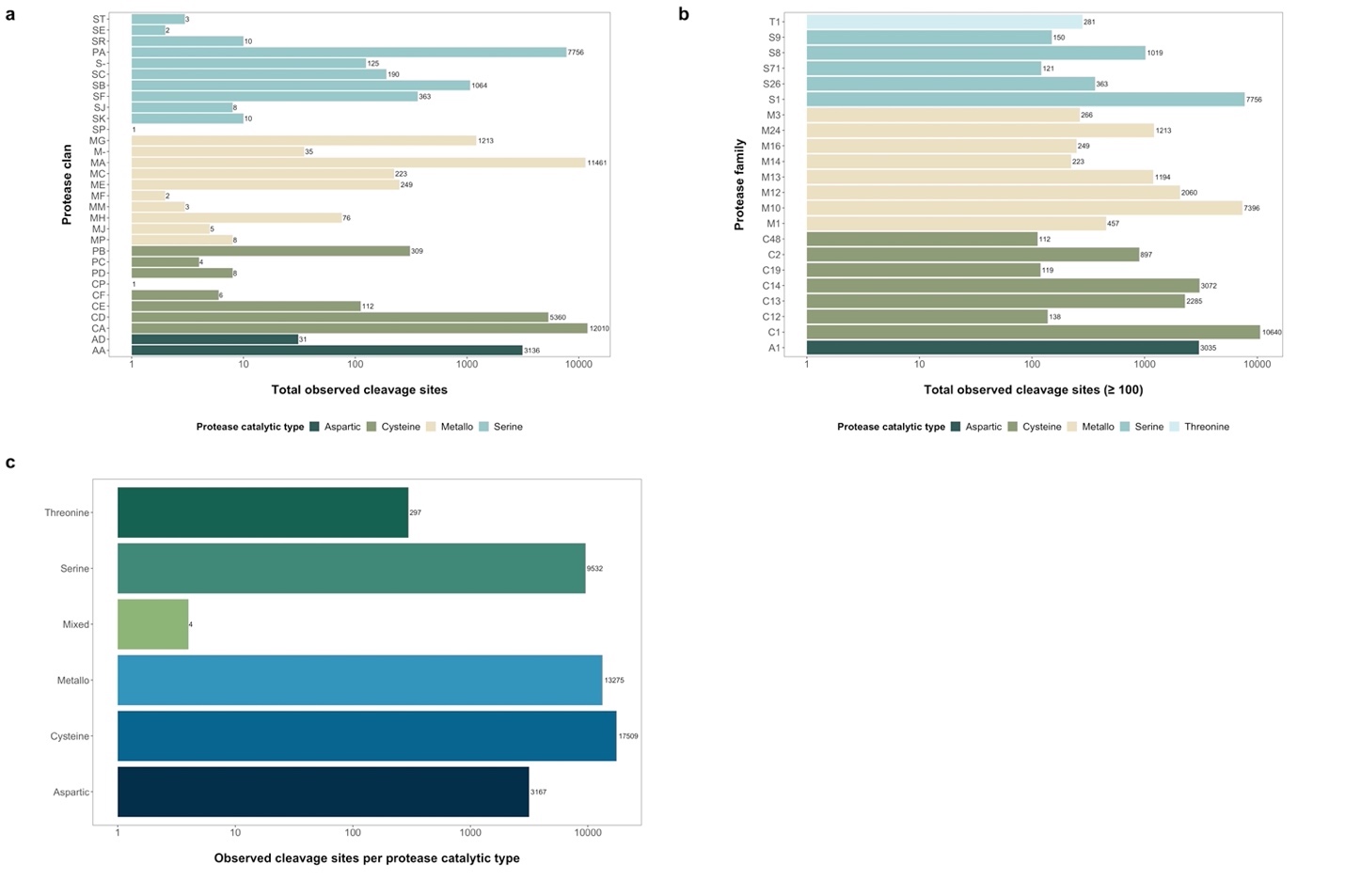

**Fig. S2. Representation of proteases among substrate cleavage sites in panCleave training and testing data (*n* = 24,817).** Plots represent total cleavage sites arranged by **(a)** protease clan, **(b)** family, and **(c)** catalytic type, as defined by the MEROPS Peptidase Database (1).

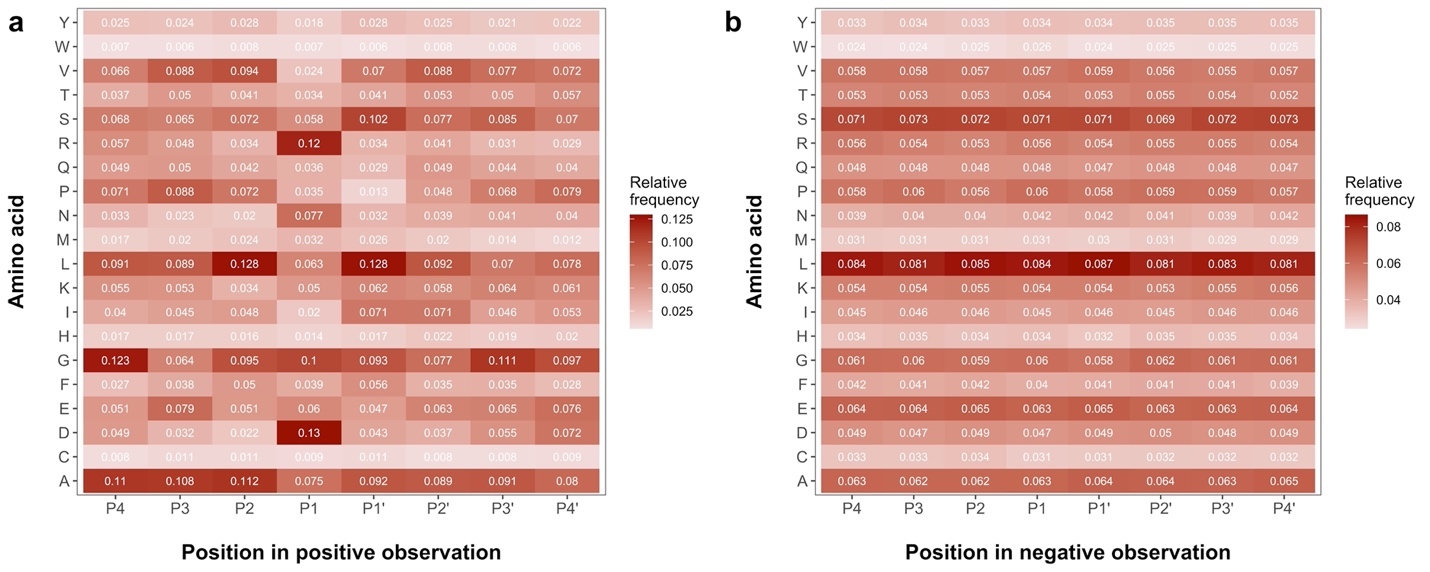

**Fig. S3. Amino acid frequencies by residue position for all training and testing data.** Amino acid relative frequencies are reported by residue position in **(a)** 8-residue positive observations (*n* = 24,817) and **(b)** 8-residue negative observations (*n* = 24,817). Cleavage takes place between positions P1 and P1’ in positive observations.

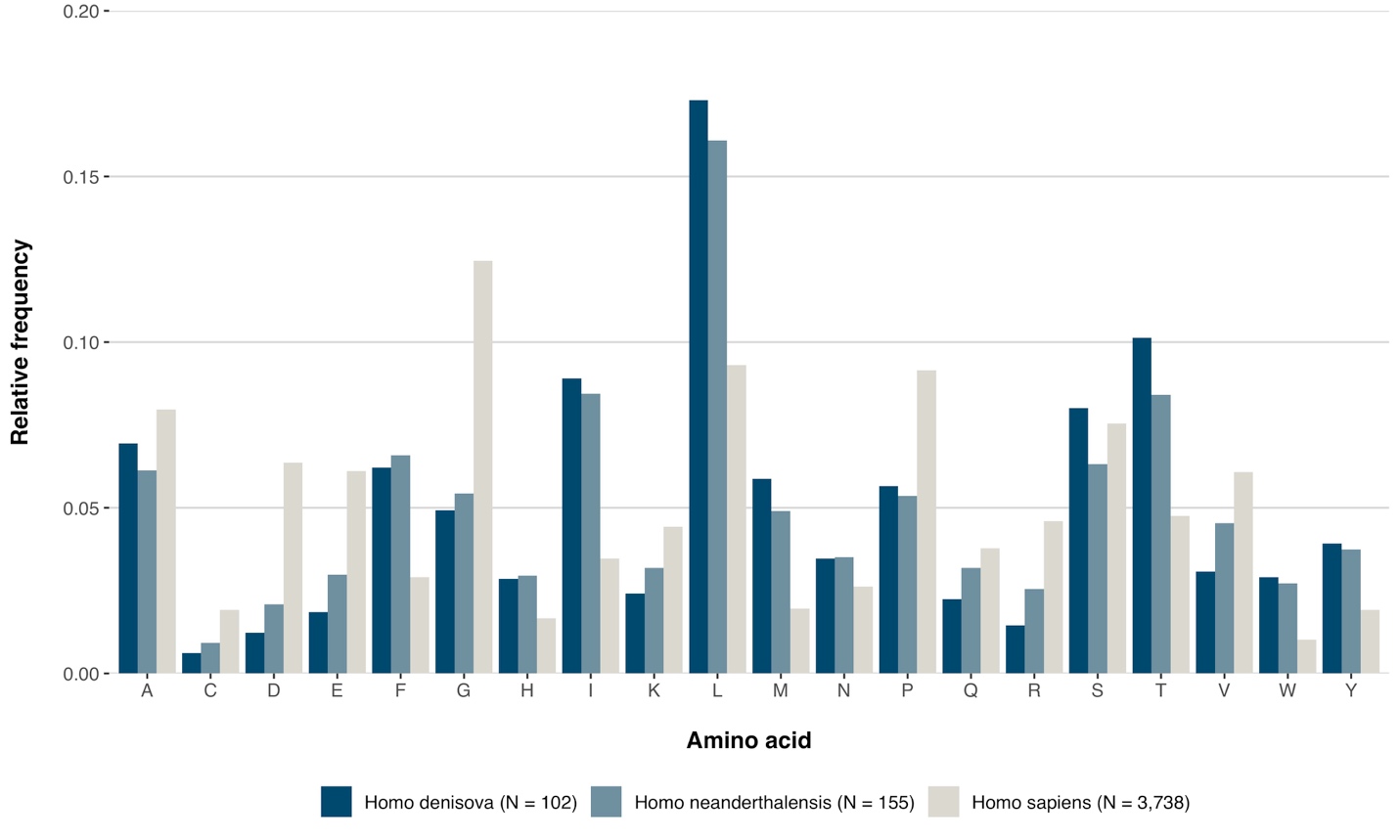

Fig. S4. Relative frequencies of amino acids for all unique panCleave fragments per taxon. This plot includes all generated fragments, not just those filtered for synthesis or with demonstrated activity.

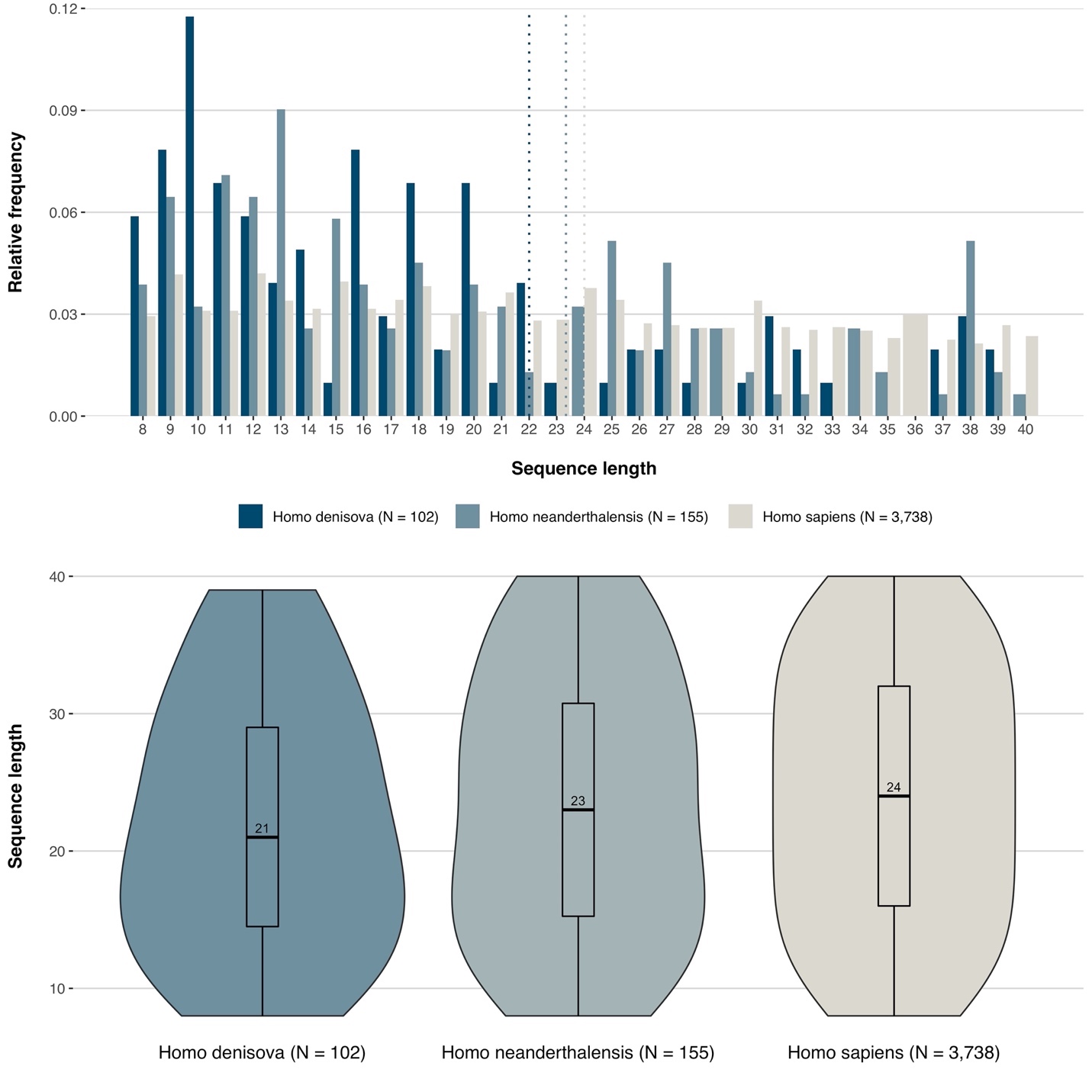

**Fig. S5. Fragment length distributions for all unique panCleave fragments per taxon.** This plot includes all generated fragments, not just those filtered for synthesis or with demonstrated activity.

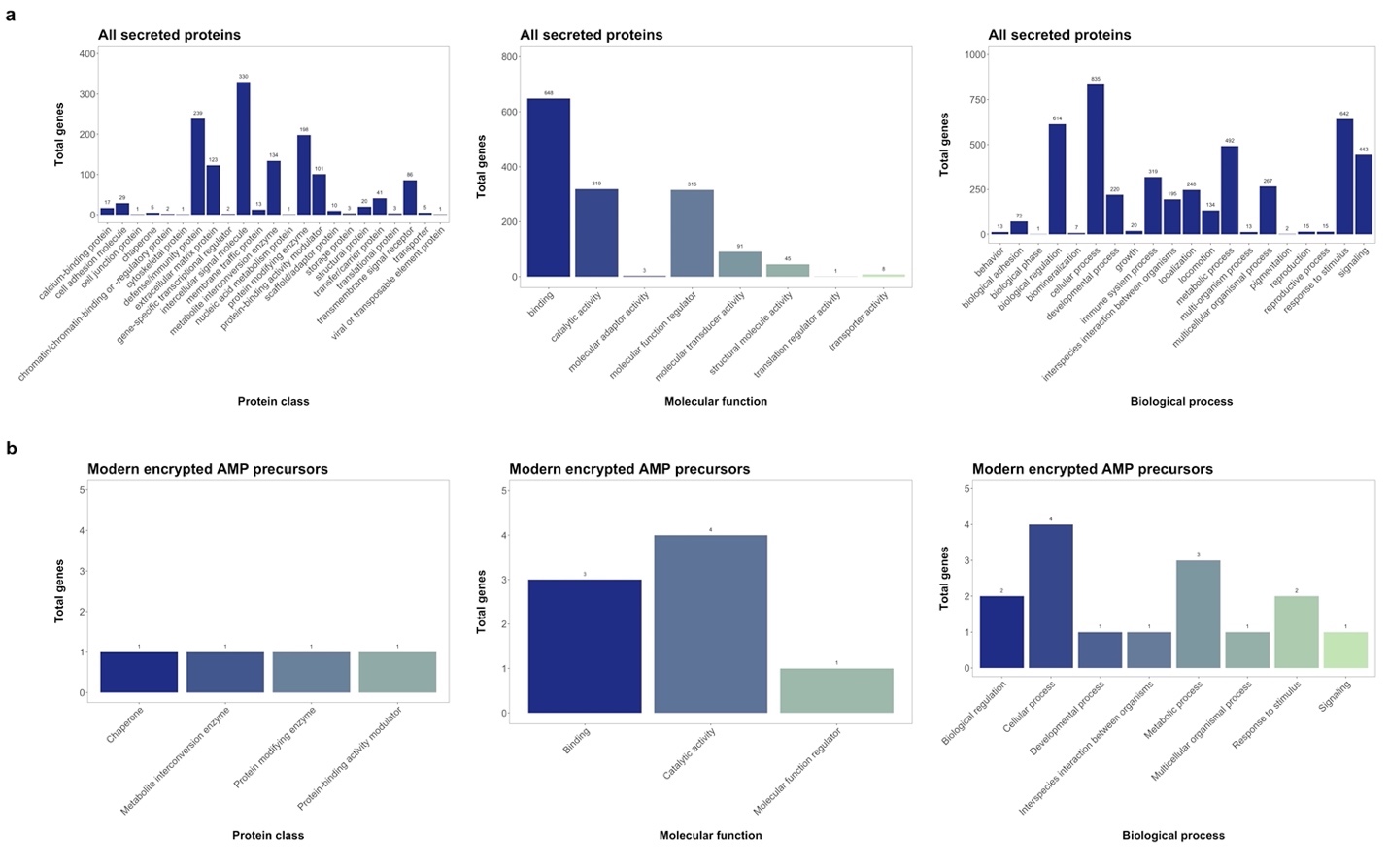

**Fig. S6. Protein classes, molecular functions, and biological processes represented by all queried human secreted proteins and by precursors of modern encrypted peptides discovered in the present work.** (**a**) All secreted proteins available in UniProt (*29*). (**b**) The precursor proteins for all modern encrypted peptides with antimicrobial activity. Data were obtained from PANTHER (<http://www.pantherdb.org/>) (*31*).

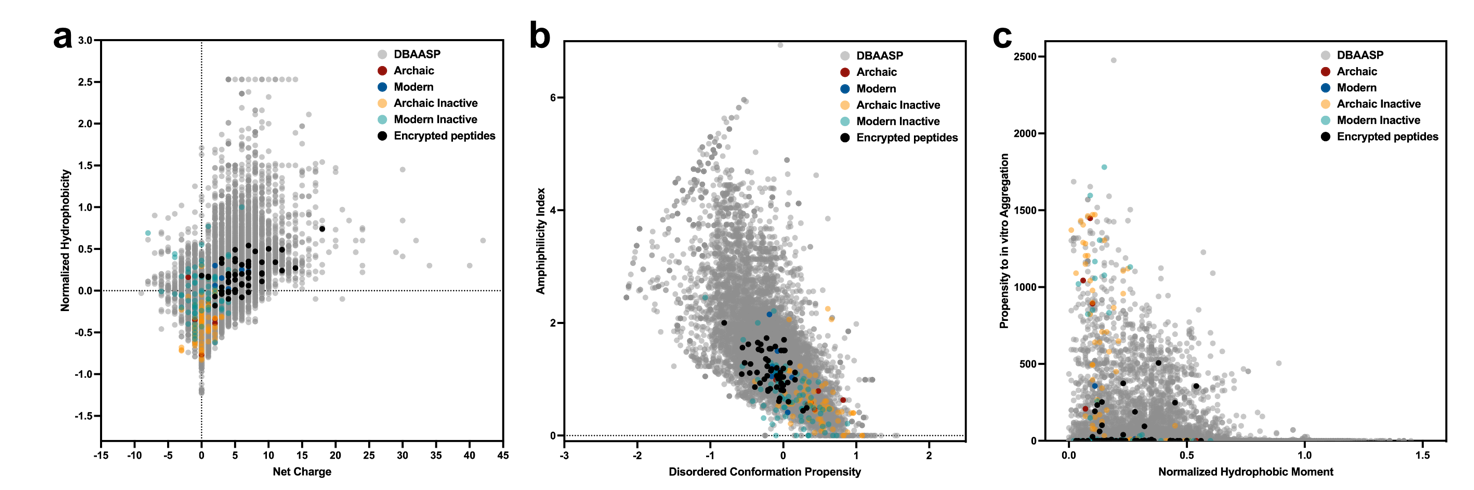

**Fig. S7. Physicochemical features of encrypted peptides identified by panCleave compared to known AMPs and other encrypted peptides.** **(a)** Net charge vs. hydrophobicity normalized according to the length of the peptide. Net charge directly influences the initial electrostatic interactions between the peptide and negatively charged bacterial membranes, and hydrophobicity directly influences the interactions of the peptide with lipids in the membrane bilayers. **(b)** Amphiphilicity index vs. disordered conformation propensity; both properties closely correlated with AMP mechanism of action. **(c)** Propensity to aggregate *in vitro* vs. hydrophobic moment normalized by peptide length; propensity to aggregate correlates with AMP toxicity.

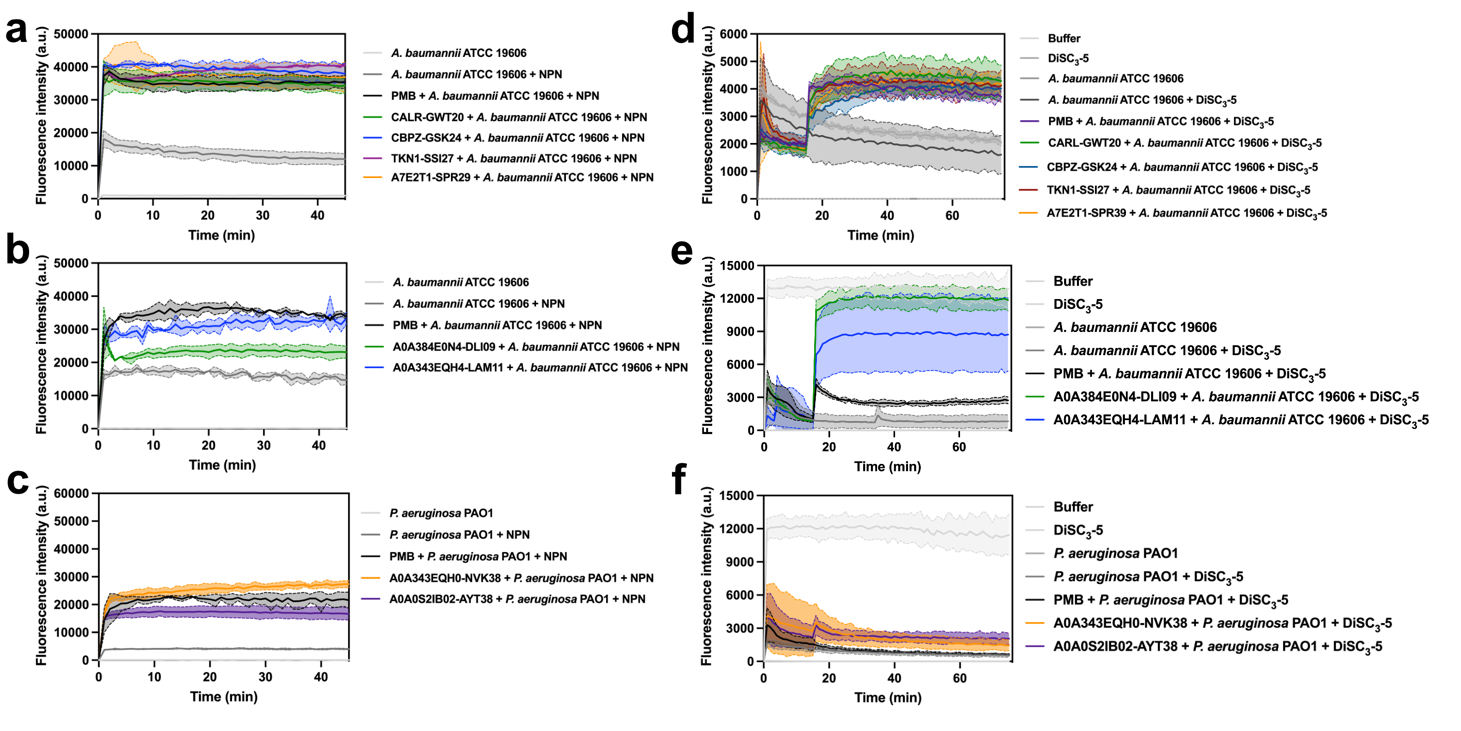

**Fig. S8. Mechanism of action of modern and archaic encrypted peptides.** Permeabilization assays with the fluorescent probe 1-(N-phenylamino)naphthalene (NPN); effect of **(a)** modern encrypted peptides and **(b)** archaic encrypted peptides on against *A. baumannii* cells, and **(c)** archaic encrypted peptides on *P. aeruginosa* PA01 cells. Depolarization assays with the hydrophobic probe 3,3′-dipropylthiadicarbocyanine iodide [DiSC_3_-(5)]; effects of **(d)** modern encrypted peptides and **(e)** archaic encrypted peptides on *A. baumannii* cells, and **(f)** archaic encrypted peptides on *P. aeruginosa* PA01 cells. All panels show the raw fluorescence intensity data obtained in the experiments.

**Fig. S9. Outer membrane permeabilization and cytoplasmic membrane depolarization triggered by archaic encrypted peptides.** Relative fluorescence values of archaic encrypted peptides compared to the untreated control**.** **(a)** Permeabilization of the outer membrane using the probe 1-(N-phenylamino)naphthalene (NPN) and **(b)** depolarization of the cytoplasmic membrane indicated by the probe 3,3′-dipropylthiadicarbocyanine iodide [DiSC_3_-(5)] of *P. aeruginosa* PA01 cells. Both peptides depolarized membranes more strongly than polymyxin B (control). A0A343EQH0-NVK38 permeabilized outer membranes more strongly than PMB or A0A0S2IB02-AYT38.
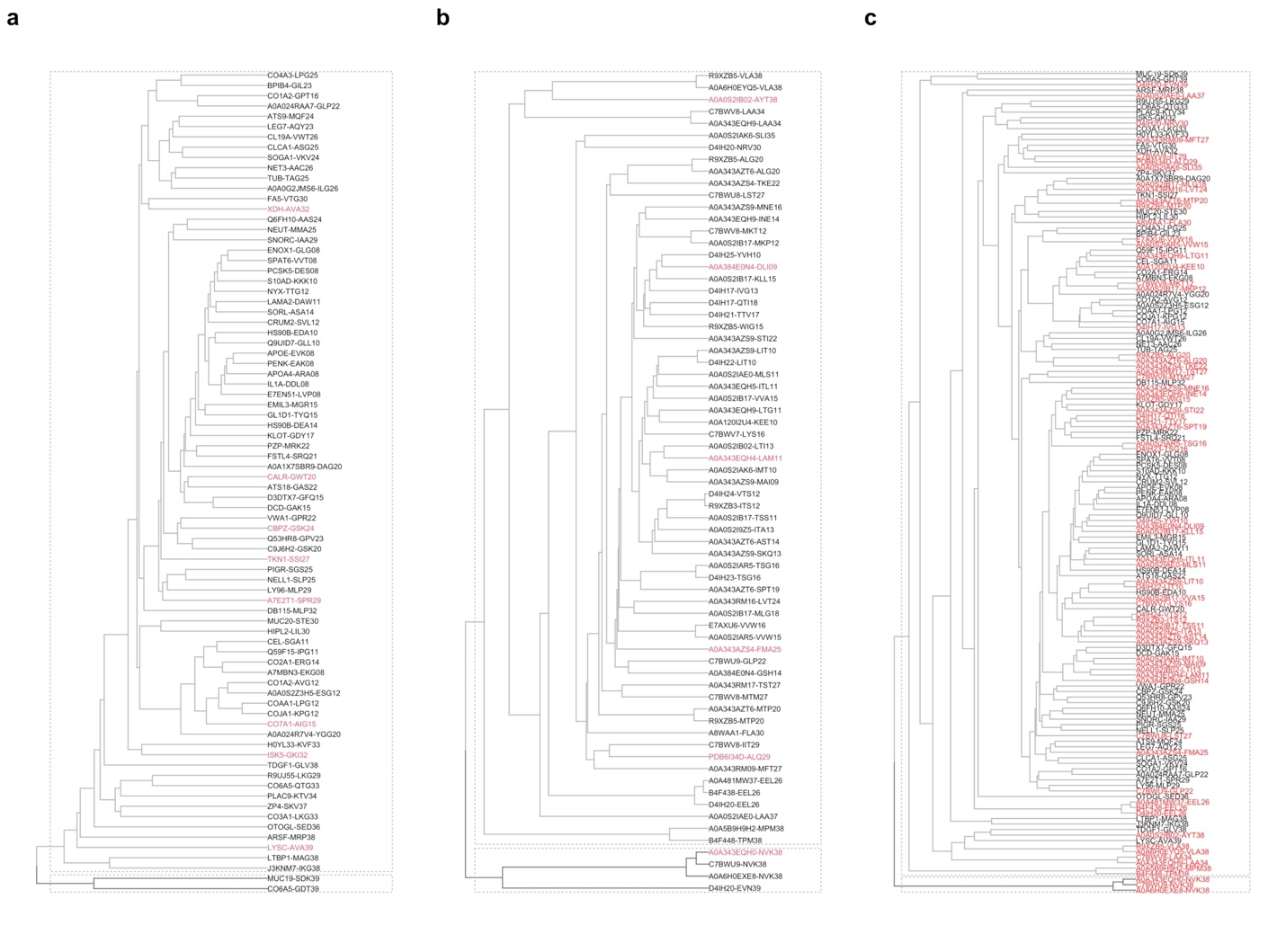

**Fig. S10. Hierarchical *k*-means clustering of synthesized peptides.** Archaic (**a**) and modern (**b**) encrypted peptides did not cluster neatly according to the presence (red labels) or absence (black labels) of antimicrobial activity. Likewise, hierarchical *k*-means clustering of archaic (red labels) and modern (black labels) in panel (**c**) did not reveal clean separation of archaic and modern fragments. Values for *k* (*i.e.*, total clusters; *k* = 2 for all subfigures) were selected based on joint evaluation of gap statistic, average silhouette score, and within-cluster sum of squares methods. Data were represented using the ProtFP encoding method (*2*) and scaled prior to clustering.
